# The hagfish genome and the evolution of vertebrates

**DOI:** 10.1101/2023.04.17.537254

**Authors:** Ferdinand Marlétaz, Nataliya Timoshevskaya, Vladimir Timoshevskiy, Oleg Simakov, Elise Parey, Daria Gavriouchkina, Masakazu Suzuki, Kaoru Kubokawa, Sydney Brenner, Jeramiah Smith, Daniel S. Rokhsar

## Abstract

As the only surviving lineages of jawless fishes, hagfishes and lampreys provide a critical window into early vertebrate evolution. Here, we investigate the complex history, timing, and functional role of genome-wide duplications in vertebrates in the light of a chromosome-scale genome of the brown hagfish *Eptatretus atami*. Using robust chromosome-scale (paralogon-based) phylogenetic methods, we confirm the monophyly of cyclostomes, document an auto-tetraploidization (1R_V_) that predated the origin of crown group vertebrates ∼517 Mya, and establish the timing of subsequent independent duplications in the gnathostome and cyclostome lineages. Some 1R_V_ gene duplications can be linked to key vertebrate innovations, suggesting that this early genomewide event contributed to the emergence of pan-vertebrate features such as neural crest. The hagfish karyotype is derived by numerous fusions relative to the ancestral cyclostome arrangement preserved by lampreys. These genomic changes were accompanied by the loss of genes essential for organ systems (eyes, osteoclast) that are absent in hagfish, accounting in part for the simplification of the hagfish body plan; other gene family expansions account for hagfishes’ capacity to produce slime. Finally, we characterise programmed DNA elimination in somatic cells of hagfish, identifying protein-coding and repetitive elements that are deleted during development. As in lampreys, the elimination of these genes provides a mechanism for resolving genetic conflict between soma and germline by repressing germline/pluripotency functions. Reconstruction of the early genomic history of vertebrates provides a framework for further exploration of vertebrate novelties.

## Introduction

Hagfishes are deep-sea jawless vertebrates displaying a scavenger lifestyle and a prodigious capacity to produce mucus (Spitzer and Koch, 1998) (**Figure 1a**). Since hagfishes lack several key characters shared by lampreys and gnathostomes such as definitive vertebrae (Ota et al., 2011), lensed eyes with oculomotor control, and electroreceptive sensory organs (Miyashita et al., 2019; Shimeld and Donoghue, 2012), cladistic morphological analysis suggested that the hagfishes diverged before the split between lampreys and jawed vertebrates (the ‘Craniata’ hypothesis, **Figure 1b**)(Janvier, 1981). Yet, both hagfishes and lampreys stand apart from other vertebrates based on the absence of jaws and other shared gnathostome characters such as bone and dentine (Janvier, 2015) leading others to group them together as cyclostomes (the ‘Cyclostomata’ hypothesis, **Figure 1b**) (Duméril, 1812). Molecular phylogenies generally favour cyclostome monophyly, implying that hagfishes are secondarily simplified (Delsuc et al., 2006; Kuraku and Kuratani, 2006). Under either the Craniata or Cyclostomata hypothesis, living hagfishes provide a critical perspective on the evolutionary history of the vertebrate lineage, reflecting ancestral features of vertebrates or evolutionary changes within the hagfish lineage, respectively.

**Figure 1.**
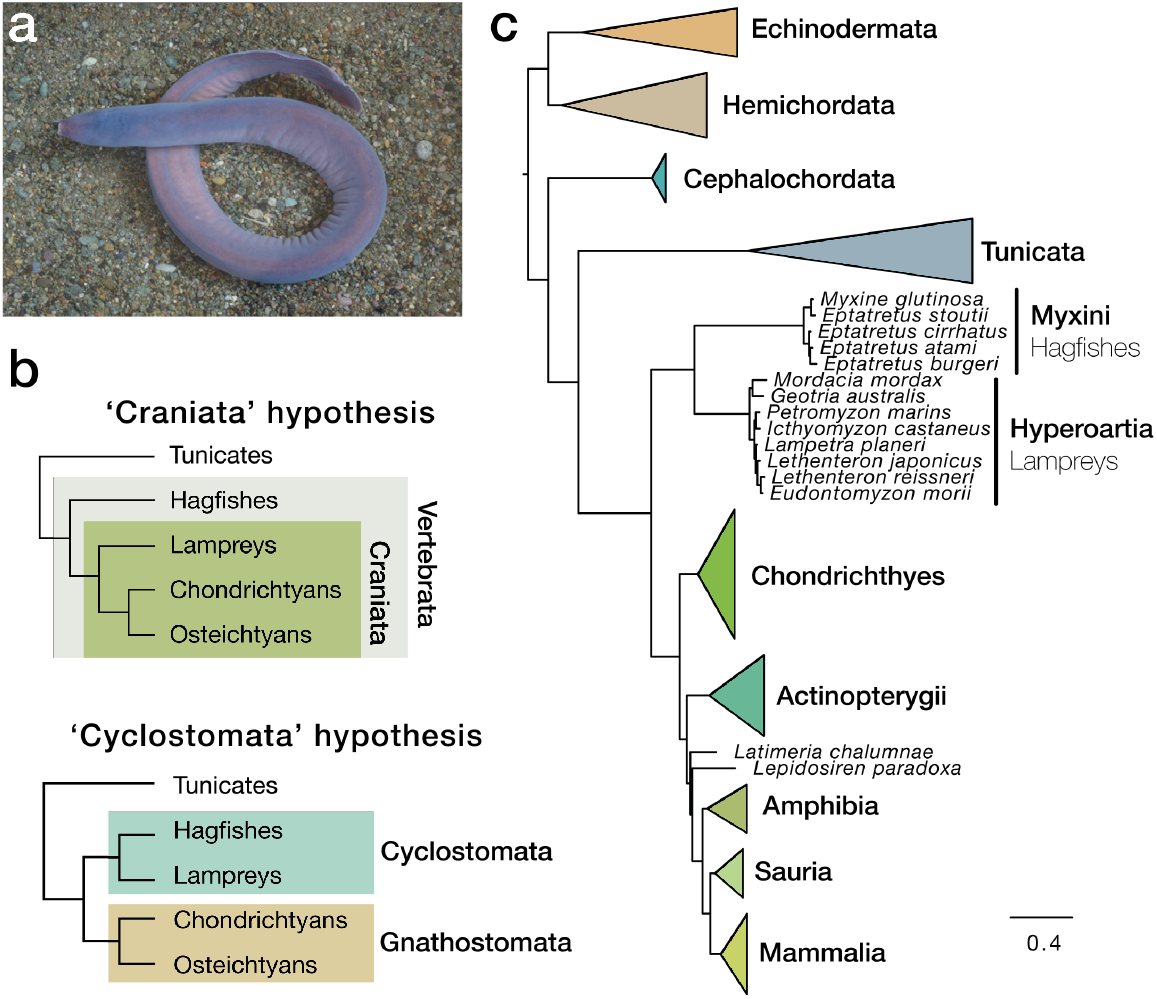
Phylogenetic position of hagfish and multigene support for the cyclostome hypothesis. a, Picture of a brown hagfish, *Eptatretus atami* (photo credit K. Kubokawa). **b**, Distinct hypothesis regarding early vertebrate evolution. **c**, Phylogenetic reconstruction of deuterostome relationships using 176 genes and 61,939 positions selected as least saturated using a site-heterogeneous model (CAT+GTR). topology is robust to composition heterogeneity and similar to what was obtained for all 1,467 genes using site-homogeneous models (**Figure S3**).

The sequence and timing of the genome duplication events that took place in early vertebrate evolution, as well as their impact on genomic architecture, is a crucial and controversial question to understand vertebrate origins which remain elusive and controversial (Kuraku, 2008; Kuraku et al., 2009a). Comparisons among chromosome-scale assemblies of amphioxus, lamprey and gnathostome genomes have provided some key insights (Nakatani et al., 2021; Simakov et al., 2020; Smith et al., 2018). Analyses of lamprey genome sequences as well as several gene-centric studies have suggested that cyclostomes share at least one of the two rounds of whole genome duplication (2R) that took place at the origin of vertebrates (Escriva et al., 2002; Furlong and Holland, 2002; Kuraku, 2008; Nakatani et al., 2021; Smith et al., 2013; Smith and Keinath, 2015). Comparisons with the (unduplicated) chromosome-scale genome of amphioxus enabled the reconstruction of the extensive chromosomal fusions and rearrangements that followed the 2R in the gnathostome lineage (Simakov et al., 2020). Hox gene clusters have historically served as markers for deciphering whole genome duplication history, and six hox clusters have been identified in lamprey and hagfish, but no clear pairwise orthology relationships could be established among gnathostome and cyclostome Hox genes, indicative of complex or independent duplication histories (Mehta et al., 2013; Pascual-Anaya et al., 2018). The perspective from the hagfish genome, the other cyclostomes lineage, is therefore essential to delineate the sequence of events that led to the genomic organisation of modern vertebrate lineages.

Unlike most other animals (Drotos et al., 2022; Smith et al., 2021) the large-scale structure of lamprey and hagfish genomes are modified during (normal) development by programmed elimination of a portion of their genome (Nakai et al., 1991; Smith et al., 2018, 2009). While hagfish were the first vertebrate species to be recognized as undergoing developmentally programmed elimination of chromatin (Nakai et al., 1991), our understanding of the content of eliminated hagfish chromosomes is thus far limited to the characterization of several satellite repeats that are highly enriched on these chromosomes (Goto et al., 1998; Kojima et al., 2010; Kubota et al., 1993; Nabeyama et al., 2000). In lampreys, the comparison of the germline and somatic genome sequences revealed that germline chromosomes contain large numbers of genes with putative functions in the germline (Smith et al., 2018, 2009). The extent to which hagfish eliminates genes during development remains unresolved, and correspondingly, it is unknown whether DNA elimination might reflect an ancestral, shared-derived or independently-evolved trait in the cyclostomes.

Here, we report a chromosome-scale assembly of a hagfish genome and use this assembly to resolve the gene content of retained and eliminated portions of the genome. This chromosomal sequence allows analyses that leverage both molecular phylogeny and conserved synteny to reconstruct duplication and divergence events that shaped the genomes of ancestral vertebrate, cyclostome and gnathostome lineages, as well as their impact on the emergence of genes involved in the evolution of vertebrate characters.

### The derived chromosomal organisation of hagfish

We sequenced the germline genome of the brown hagfish *Eptatretus atami* (formerly *Paramyxine atami*) using a combination of short and long reads from testes and organised the assembly into chromosomes using proximity ligation data from somatic tissue (**Table S1**). Our *E. atami* assembly spans 2.52 Gb and includes 17 large chromosomal scaffolds, consistent with the expected somatic karyotype (2n = 34) (**Figure S1a** and **Table S2**). The length of the assembly is intermediate between the fluorescence-based estimates of genome size for somatic (2.01 Gb) and germline (3.37 Gb) cells (Nakai et al., 1995, 1991), consistent with k-mer estimates (2.02 and 3.28 Gb, respectively, **Figure S1b, Methods**). The *E. atami* germline genome also includes seven highly repetitive chromosomes that are completely eliminated during development, and whose sequence is present in our assembly as shorter fragments, as similarly found for the highly repetitive germline-specific chromosomes of lampreys (Smith et al., 2018; Timoshevskaya et al., 2023) and songbirds(Kinsella et al., 2019). We annotated 28,469 genes of which 22,663 show similarity with the protein-coding complement of another species (**Supp. File 1**).

The karyotype of hagfish differs from that of lampreys, which have 2n∼168 small somatic chromosomes plus additional germline-specific chromosomes (Timoshevskiy et al., 2019). Despite their distinctive genomic organisation, hagfish and lamprey chromosomes are directly related: each hagfish chromosome corresponds to a fusion of 2 to 6 lamprey chromosomes

(**Figure 2b**). Conversely, each lamprey chromosome is typically associated with only one hagfish chromosome. While chromosomal identity and colinear gene order are surprisingly highly conserved between the lampreys *Petromyzon marinus* and *Lethenteron reissneri* that diverged ∼162 Ma ago (**Figure S2a**), collinearity is not preserved between lamprey and hagfish (**Figure S2b**). Hagfish and lamprey also have distinctive repetitive landscapes **(Figure S1c**,**d**).

**Figure 2.**
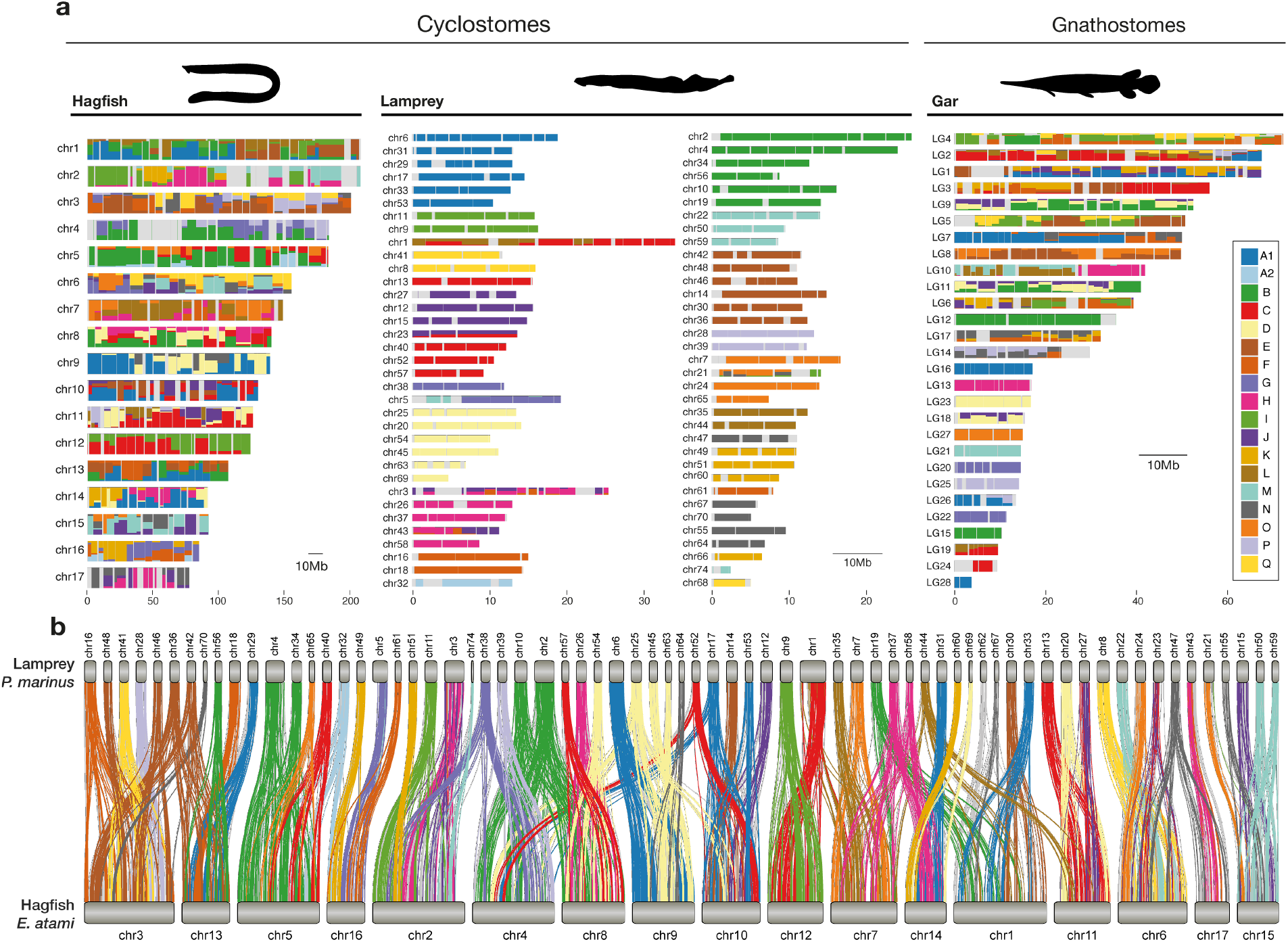
Genomic and syntenic architecture in cyclostomes and vertebrates. **a**, Distribution of genes derived from the 18 ancestral chordate linkage groups (CLGs) in the chromosomes of hagfish, lamprey and gar chromosomes. Bins contain 20 genes and only CLGs from chromosomes showing reciprocal enrichment are plotted (fisher’s test). **b**, Syntenic relationship between lamprey and hagfish chromosomes showing that hagfish chromosomes are typically fusions of multiple lamprey chromosomes. Each line corresponds to a single-copy orthologue labelled by CLGs. CLG colours are the same as in **a**.

Using the cephalochordate amphioxus as an outgroup, the ancestry of hagfish, lamprey, and other vertebrate chromosomes (or chromosomal segments) can be traced back to the ancestral proto-vertebrate (chordate) linkage groups (CLGs) (Simakov et al., 2020). The majority of lamprey chromosomes derive from a single CLG, with each CLG giving rise to 2 to 6 lamprey chromosomes due to subsequent genome duplications (see below) (**Figure 2a)**. The simple correspondence between lamprey chromosomes and ancestral chordate linkage groups(Nakatani et al., 2021; Simakov et al., 2020; Smith and Keinath, 2015) implies that the chromosomal organisation of the hagfish-lamprey ancestor more closely resembled contemporary lamprey. In contrast, hagfish chromosomes are derived by the irreversible process of fusion-with-mixing of lamprey-like ancestral chromosomes, analogous to but distinct from the fusion-with-mixing events found in gnathostomes (Simakov et al., 2020) (**Figure 2b**).

### Phylogenomics confirm the monophyly of cyclostomes

The chromosome-scale hagfish genome assembly is a valuable resource for addressing the craniate-vs-cyclostome debate. Previous molecular phylogenies (Delsuc et al., 2006; Kuraku and Kuratani, 2006) as well as analysis of vertebrate miRNAs (Heimberg et al., 2010) (but see (Thomson et al., 2014)) supported cyclostomes, but these early studies were constrained by limited gene and taxonomic sampling (Kapli et al., 2020). Moreover, lampreys and hagfish show both nucleotide and amino-acid composition bias (Kuraku and Kuratani, 2006) and their complex history of genome duplication and gene loss complicates the identification of orthologs (Furlong and Holland, 2004).

We found robust support for cyclostome monophyly using a set of 1,553 orthologous genes that was inferred using hagfish and lamprey genome data as well as new transcriptome sequences for the hagfish *Myxine glutinosa* (**Table S3**). Cyclostome monophyly is recovered with a partitioned analysis (**Figure S3a**), site-heterogeneous models (**Figure 1c**), and six-category amino-acid recoding to alleviate compositional bias (**Figure S3c**). Using fossil calibrations (**Table S4**), we estimated that the divergence between lamprey and hagfish took place in the Late Ordovician ∼449 Ma, with the split between cyclostomes and gnathostomes occurring close to the Cambrian-Ordovician boundary ∼493 Ma, consistent with previous estimates (Kuraku et al., 2009b). The diversification times of modern lampreys ∼182 Ma (Early Jurassic) and hagfishes ∼130 Ma (Early Cretaceous) (**Figure S3d**) are compatible with the recent discovery of a late-Cretaceous hagfish (Miyashita et al., 2019), and points to long stem lineages for both taxa.

### Genome duplications in early vertebrate evolution

The sequence and timing of the whole genome duplications that occurred early in vertebrate evolution remain controversial (Kuraku, 2008; Kuraku et al., 2009a; Nakatani et al., 2021; Sacerdot et al., 2018; Simakov et al., 2020; Smith et al., 2018; Smith and Keinath, 2015). Major unresolved questions include whether zero, one, or two whole genome duplications occurred on the vertebrate stem lineage prior to the cyclostome-gnathostome split, and the timing and nature of any subsequent lineage-specific events (**Figure 3a**). Definitive phylogenetic resolution of these questions has been frustrated by uneven rates of evolution, differential gene losses and associated ‘hidden paralogy’ (Holland et al., 2017; Kuraku, 2010; Kuraku et al., 2009a), and possible delayed rediploidization (Furlong and Holland, 2002).

**Figure 3.**
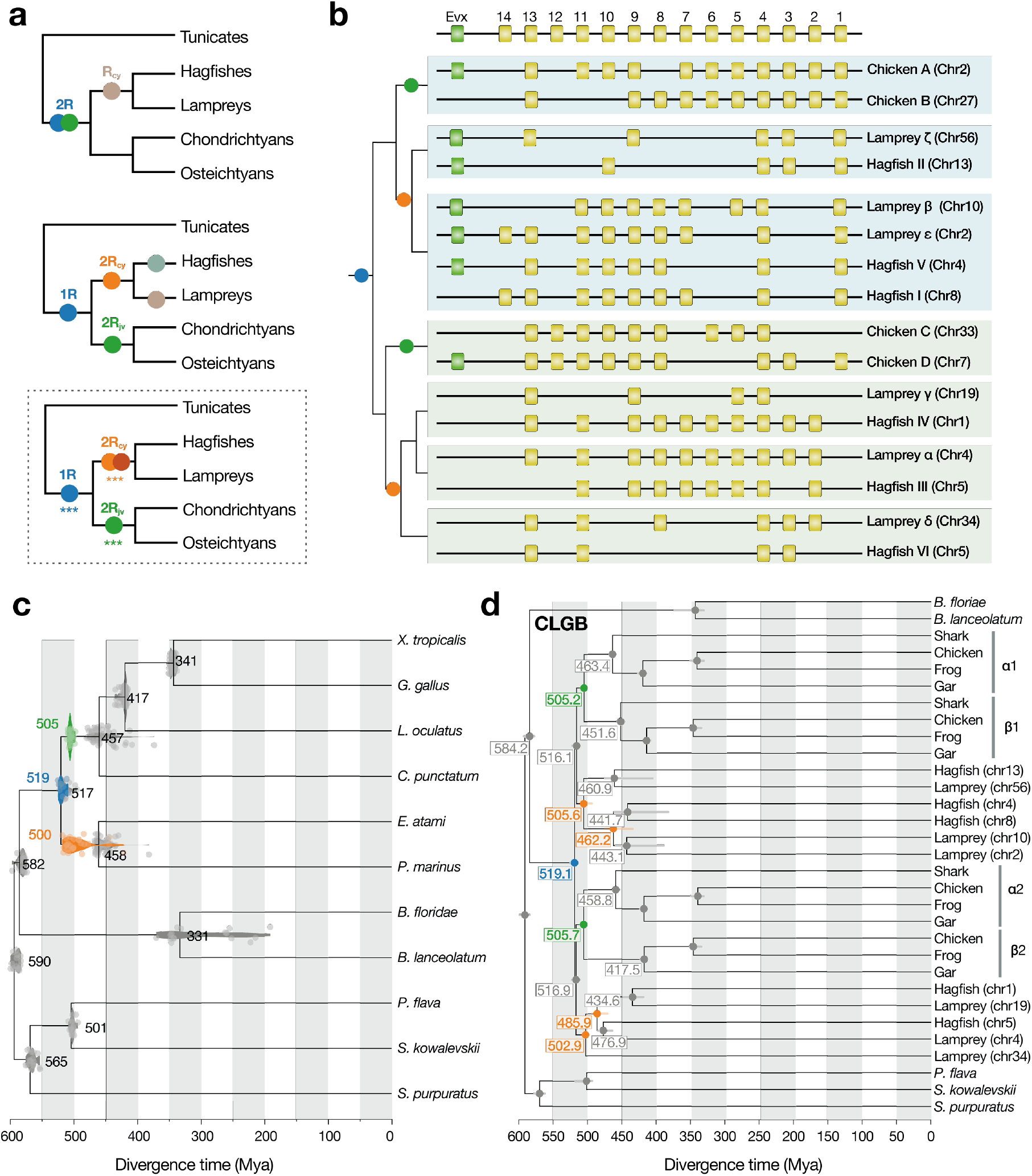
History of genome duplications in vertebrates. **a**, Alternative scenarios for whole genome duplications during early vertebrate evolution tested using WHALE with the scenario and duplication nodes receiving statistical support highlighted (see **Figure S4**). **b**, Evolutionary history of vertebrate Hox gene clusters determined by paralogon phylogeny (see also panel **d**). **c**, Timetree of deuterostomes combining results from paralogon-based molecular dating analyses. Distribution of divergence times for speciation (grey) and duplication (coloured as in **a**) nodes inferred for the 17 CLGs is shown at each node (**Table S6**). Each node is labelled with the median divergence time across CLGs and individual dates for distinct CLGs are plotted as dots with as an overlay the distribution of inferred divergence time for all CLGs. **d**, Example of topology and divergence times obtained using segments derived from the Hox bearing CLG B. Species and datasets used are listed in **Table S5**. Vertebrate species: *X. tropicalis*, western clawed frog; *G. gallus*, chicken; *L. oculeatus*, spotted gar; *C. punctatus*, brownbanded bamboo shark. Outgroups include two amphioxus (*B. floridae, B. lanceolatum*), hemichordates (*S. kowalevskii* and *P. flava*), and one echinoderm (*S. purpuratus*).

Recently, we proposed a scenario for early vertebrate evolution in which an initial auto-tetraploidization, likely shared by both gnathostomes and cyclostomes, and was followed by a gnathostome-specific allo-tetraploidization (Simakov et al., 2020). Additional whole genome duplications (Simakov et al., 2020) or a hexaploidization (Nakatani et al., 2021) on the lamprey lineage have been suggested, but whether these later events inferred from the lamprey genome occurred before, after, or even contemporaneously with the lamprey-hagfish split has not been investigated for lack of a chromosome-scale hagfish genome (**Figure 3a**).

We first tested different vertebrate whole genome duplication scenarios using WHALE (Zwaenepoel and Van de Peer, 2019), a method based on probabilistic reconciliation of gene and species trees (**Table S5**, see also **Methods**). We recovered significant support for the scenario placing the 1R on the ancestral lineage that gave rise to all extant vertebrates (1R_V_), followed by clade-specific genome-wide duplications in the gnathostomes (2R_JV_) and cyclostomes (2R_CY_) lineages (all Bayes Factors BF_Null_vs_WGD_ < 10^−3^) **(Figure S4**). In contrast, we found no support for lineage-specific genome-wide duplications in lamprey or hagfish **(Figure S4**), consistent with their near 1:1 segmental relationship (**Figure 2b**).

We confirmed the order and timing of early vertebrate whole genome duplications with a complementary new approach leveraging deeply conserved synteny (Simakov et al., 2022, 2020). We reconstructed molecular phylogenies from concatenated blocks of chromosomally linked genes that derive from each of the proto-vertebrate chromosomes (the ancestral chordate linkage groups CLGs of ref. (Simakov et al., 2020)) (**Figure 3c,d** and **Table S6**). Such descendants of the ancestral chordate chromosomes are called ‘paralogons’ (Coulier et al., 2000). Since the genes of each paralogon are expected to have experienced the same evolutionary history (barring homoeologous recombination), the problem of ‘hidden paralogy’ is avoided; combining linked genes also increases the number of phylogenetically informative characters relative to individual gene trees (**Table S6**). We estimated paralogon phylogenies and timings of divergences and duplications independently for each ancestral proto-vertebrate chromosome (Simakov et al., 2020) with the hypothesis that these trees should display consistent patterns and timing of divergence and duplication.

Among gnathostomes, we find robust support for the proposal of an early auto-tetraploidization (1R_V_) followed by a later gnathostome-specific allo-tetraploidization (2R_JV_) (see, *e*.*g*., CLGB in **Figure 3d; Figure S6**; **Supp. file 2**). Based on patterns of gene loss, the gnathostome-specific 2R_JV_ was previously hypothesised to be an allo-tetraploidization that brought together genomes from two now-extinct gnathostome progenitors, alpha and beta (Simakov et al., 2020). Alpha and beta share characteristic gnathostome-specific chromosomal fusions that are absent from cyclostomes (*e*.*g*., CLG O & E or C & L, **Figure 2a**, see also (Simakov et al., 2020)). Paralogon trees resolve several ambiguities in the assignment of beta segments to their 1R alpha counterparts (Simakov et al., 2020) (**Table S7**).

We estimated that the alpha and beta progenitors of gnathostomes diverged in the mid-late Cambrian ∼505 Mya (**Figure 3b, Figure S6**) based on calibration against the fossil record (**Table S4, Methods**). These progenitors later hybridised, in association with a genome doubling, prior to the origin of the crown group gnathostomes (i.e., the split between cartilaginous and bony fishes) near the end of the Ordovician ∼457 Mya. The timing of this hybridization/doubling event cannot be determined from molecular phylogenies but we speculate that it likely occurred within 10-15My of the alpha-beta divergence based on examples from recent allo-tetraploidization events such as *Xenopus* (Session et al., 2016) and goldfish (Chen et al., 2019).

Regarding cyclostomes, paralogon trees generally identify directly orthologous hagfish and lamprey chromosome segments, as exemplified in **Figure 2b**, although some pairings have low support and some segments have insufficient representation due to extensive gene loss (**Table S6** and **Supp. file 2**). This direct segmental orthology is consistent with cyclostome monophyly and the absence of hagfish- or lamprey-specific whole genome duplications. Conversely, we find evidence for shared genome-wide duplications on the cyclostome stem, i.e., preceding the divergence of the hagfish and lamprey lineages ∼458 Mya. While the nature of the cyclostome-specific duplications is difficult to decipher, the net effect appears to be hexaploidization(Nakatani et al., 2021) without obvious patterns of differential gene retention as typically observed after allopolyploidy involving divergent progenitors and seen in the gnathostome lineage (Simakov et al., 2020).

The broad distribution of divergence times observed between homoeologous cyclostome chromosomes (∼515-460 Mya) is consistent with an extended period of ongoing diploidization (Furlong and Holland, 2002), as seen in salmonids (Gundappa et al., 2022; Lien et al., 2016). More specifically, this state of affairs parallels proposals for the origin of hexaploidy in modern sturgeon, in which stem lineages exist in multiple interfertile ploidies (Fontana et al., 2008). Under this model, gene duplications in the cyclostome lineage would have occurred in two successive bursts, the first from the formation of a (now extinct) tetraploid stem cyclostome and the second from the hybridization of tetraploid and diploid stem cyclostomes, followed by possibly extended periods of rediploidization (**Figure S6**). The near 1:1 relationship between orthologous hagfish and lamprey chromosome segments (**Figure 2b**) suggests that this process must have been largely completed by the time the crown group cyclostomes originated.

With regard to the timing of 1R, our paralogon trees and probabilistic inference of genome duplications (WHALE) both support a single shared auto-tetraploidization on the vertebrate stem (1R_V_) predating the divergence of cyclostomes and gnathostomes. The sequence of events involving first the 1R_V_ duplication and second the cyclostome-gnathostome speciation is reflected in 10 out of 14 CLGs that show strong phylogenetic resolution (bootstrap support BP>60, **Table S6, Supp. file 2**). The absence of cyclostome derivatives on one side of the 1R_V_ branch (in 5 out of 17 CLGs) can be due to loss of paralogous segments, limited phylogenetic signal, and/or delayed diploidization (Parey et al., 2022; Robertson et al., 2017).These duplication and speciation events occurred in close succession: we estimate that 1R_V_ took place ∼519 Mya and the cyclostome-gnathostome split ∼517 Mya. We note that, following Furlong and Holland (Furlong and Holland, 2002), our estimate for 1R_V_ corresponds to the cessation of homoeologous recombination after auto-tetraploidization, and the actual duplication event could have occurred earlier.

### History of vertebrate Hox clusters

Paralogon-based phylogenies can also be used to elucidate the tangled relationships among the Hox clusters of cyclostomes and gnathostomes. While gnathostomes typically have four Hox clusters (Hox A-D), hagfish and lamprey each have six clusters (Hox I-VI and Hox α-ζ, respectively)(Mehta et al., 2013; Pascual-Anaya et al., 2018; Smith et al., 2018) (**Figure 3b**). Hox clusters are located on the descendants of chordate linkage group B, and the corresponding paralogon tree indicates a deep split between (gnathostome) HoxA, D, (lamprey) Hox β, ε, ζ, and (hagfish) Hox I, II, IV on one side and Hox B, C, Hox α, γ, δ and HoxIII, IV, VI on the other, corresponding to the 1R_V_ duplication on the vertebrate stem (**Figure 3d**). Subsequent independent duplications on the cyclostome and gnathostome lineage further expanded these two paralogous early vertebrate Hox clusters. At a finer scale, we also resolve the uncertainty regarding the correspondence between the lamprey and hagfish Hox clusters, whose relationships are supported by both the paralogon tree, as well as a concatenation of Hox and surrounding genes (‘bystanders’) (**Figure S5**). However, the relationship between hagfish Hox I and V and lamprey Hox β and ε (lamprey chromosomes 2 and 10 on **Figures 2b, 3b,d** and **S5**) has been obscured by an apparent reciprocal fission-fusion event in the cyclostome lineage.

Interestingly, Evx, a homeobox gene linked to the core Hox cluster, is missing from all cyclostome clusters that diverged from Hox C/D after 1R_V_. In contrast, gnathostomes have two Evx paralogs whose duplication can be traced to 1R_V_: Evx1 (linked to HoxA) and Evx2 (linked to Hox D). The retention of both 1R_V_-derived Evx paralogues in the gnathostome lineage is notable given the importance of these genes in the development and patterning of fins and limbs across divergent gnathostome lineages(Sordino et al., 1996). This suggests that 1R_V_ duplicates acquired roles in fin bud development and patterning very early in the evolution of the gnathostome lineage, consistent with the observation of paired fin fold morphologies in early diverging galeaspids(Gai et al., 2022).

### Did neural crest arise before or after 1R_V_?

Many developmental genes retained after WGD have been instrumental to establish vertebrate innovations (Shimeld and Holland, 2000). The resolution of early vertebrate genome duplications provides a framework for assessing whether core vertebrate characters such as neural crest, placodes, and vertebrate hormone systems arose before, after, or concomitant with 1R_V_. The logical framework for such an assessment was outlined by Wada et al. (Wada and Makabe, 2006). When paralogous genes share a common (vertebrate-specific) function, it is likely that their unduplicated ancestor also carried out that function; conversely, if only one pair of paralogs has a novel vertebrate-specific function, then either (1) that paralog acquired this function after duplication, or (2) the function preceded duplication but the function was subsequently lost in the other paralog. Importantly, while individual gene trees often have low signal, our paralogon trees are more robust and allow confident assignment of gene duplications to genes to specific whole genome events.

We considered the origin of neural crest cells relative to 1R_V_ by analysing a set of 22 gene families involved in neural crest specification and migration (Martik and Bronner, 2021; Simões-Costa and Bronner, 2015) (**Figure 4a**). We find that for many of these genes, including Tfap, Sox, Ednr, Twist and Gata, paralogs on both 1R_v_ branches are involved in neural crest-related functions (**Table S8**). This indicates that these functions were likely inherited from a single pre-1R_v_ gene in the vertebrates ancestor and suggests that the neural crest likely originated before the 1R_v_ and that post-WGD subfunctionalisation played a limited role in the emergence of this cell population, contrary to other gnathostome novelties such as limbs (Minguillon et al., 2009). Notably, the history of vertebrate genome duplication (**Figure 3c, Figure S6**) allows us to draw this conclusion based on studies of jawed vertebrate neural crest and makes predictions for functions of cyclostome orthologues whose potential role in neural crest has not yet been tested.

**Figure 4.**
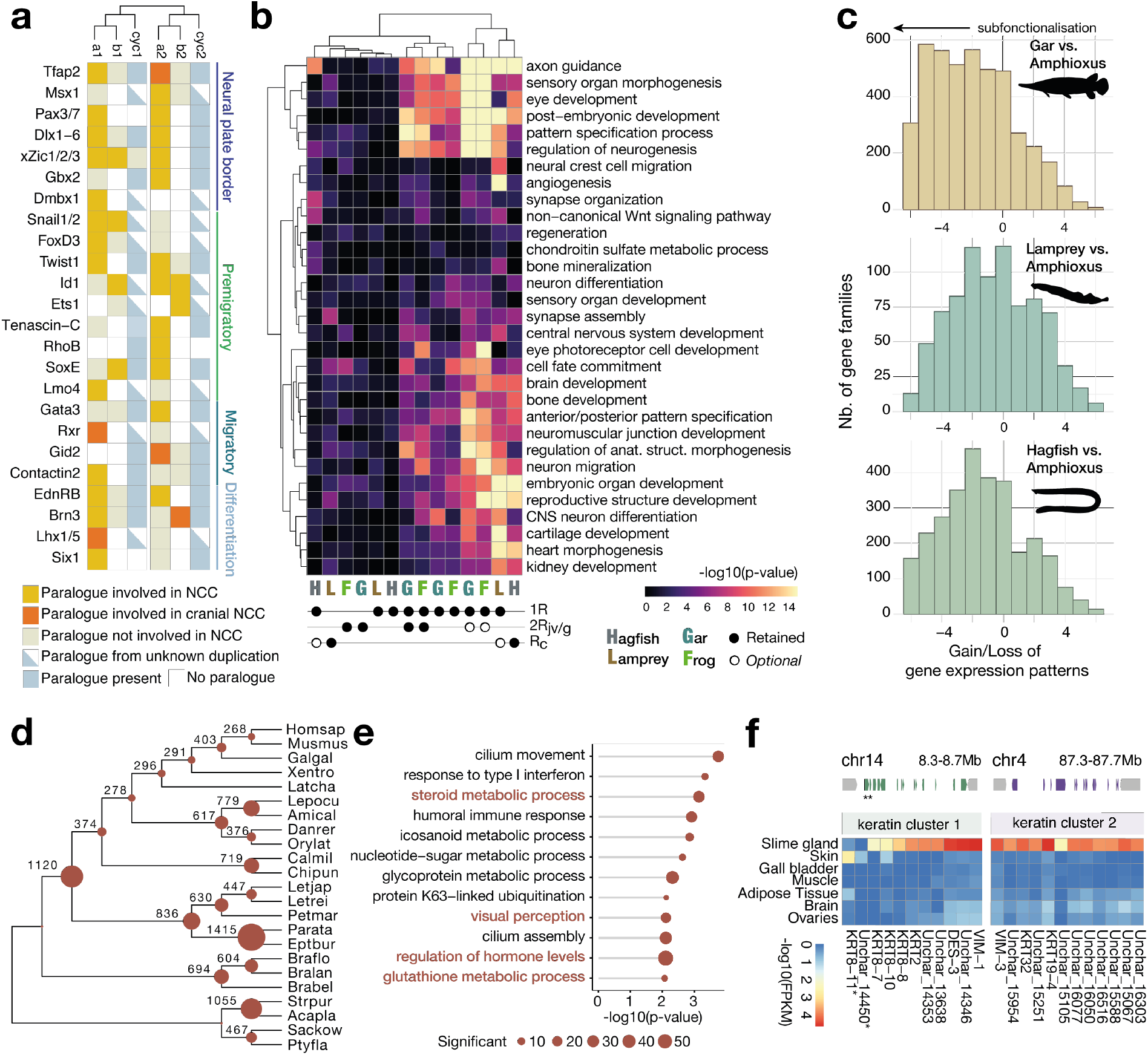
Functional impact of vertebrate WGD and gene loss in vertebrates. **a**, Key neural crest-related genes classified according to their paralogy status in regarding to the 1R_V_. **b**, Enrichment of functional annotation terms (Gene Ontology) in sets of genes showing a specific pattern of retention after vertebrate WGDs. **c**, Subfunctionalization of paralogous genes evaluated by counting expression pattern gain-and-loss. **d**, Gene family loss in deuterostomes highlighting the severe loss in the hagfish lineage. Species abbreviation is described in **Table S5. e**, Functional enrichment (GO) in lamprey for genes lost in the hagfish lineages highlighting a simplification of visual and hormonal systems. **f**, Structure and gene expression for the two clusters of alpha-keratin genes expressed in the slime gland and the skin with RNA-seq signal in blue (**Figure S8**).

While NCCs are shared by all vertebrates, the specification and patterning of the trunk and cranial NCCs appear to differ from cyclostome, osteichthyans and even amniotes, with distinct genes being involved (Martik et al., 2019). We found that genes such as Tfap and SoxE involved in the ancestral specification of both cranial and trunk NCC all seem to have paralogues on both 1R_v_ branches involved in this function. Conversely, genes such as Lhx5, Id3 or Gid2 or Dmbx that are involved in cranial NCC specification of gnathostomes do not have 1R_v_ or 2R_jv_ paralogues with a similar function, which suggests a later subfunctionalization and incorporation in the NCC GRN. Interestingly, Cadherin 6, 9 and 10 that play a role in migration of NCCs in tetrapods underwent an independent lineage-specific tandem duplication in sarcopterygians (**Supp. file 3**), which is consistent with the successive incorporation of new genes in the control of cranial NCCs during the course of gnathostome and tetrapod evolution (Martik et al., 2019). Despite the extensive gene loss experienced on the hagfish lineage (see below), we recovered homologues for most of the NCC-related genes that we investigated, but further functional studies would be necessary to determine whether subsequent 2R_Cy_ paralogues could have been incorporated in NCC-related functions. (Martik and Bronner, 2021) (**Table S8**).

### A distinct fate for paralogues in cyclostomes

Previous work revealed that paralogues retained after two rounds of gnathostome genome duplications were functionally enriched in terms associated with the regulation of development and nervous system activity (Putnam et al., 2008). To determine whether these genes were retained in a similar fashion in the cyclostome lineage, preferentially in 1R_V_ or subsequently after lineage-specific 2R, we conducted functional enrichment tests on paralogue sets showing distinct retention patterns (**Figure 4b**). We recovered the terms previously found enriched in gnathostome paralogues (e.g., axon guidance, embryonic organ development), but interestingly, we found them preferentially associated with retention of paralogs arising from the pan-vertebrate 1R_V_ but not the gnathostome-specific 2R_JV_ **(Figure 4b**). The similar pattern of functional enrichment in lamprey, gar, and frog is consistent with the occurrence of 1R_V_ before the cyclostome-gnathostome split. In contrast, hagfish has lost some 1R_V_ paralogues that persist in lamprey. These 1R_V_-derived paralogues were therefore initially retained in the cyclostome lineage but subsequently lost in hagfish (e.g. embryonic organ development) **(Figure 4b**). Such increased gene losses and functional divergence in hagfish could account for its derived morphology.

The fate of paralogues after WGD is often related to their acquisition of more specific expression domains that can explain subfunctionalization and functional innovation (Lynch and Conery, 2000; Marlétaz et al., 2018). To examine patterns of expression divergence in gnathostomes and cyclostomes, we compared gene expression of paralogues in a consistent set of six organs in amphioxus, gar, lamprey, and hagfish. Comparing a set of 3,009 gene families in which paralogues were retained, we observed that the gar displays a higher level of gene expression specificity than lamprey and hagfish, with the hagfish showing the least specificity (**Figure S7b**). We further examined the number of expression patterns gained or lost in the same gene family between amphioxus and the other species, which confirms a lower level of subfunctionalization in cyclostomes than in gnathostomes (**Figure 4b**). Finally, we used gene expression clustering (WGCNA) to ask whether specific organs show significant enrichment of paralogous genes **Figure S8**). Interestingly, we found that only neural tissue displays such a pattern of enrichment in both gnathostomes (gar) and in hagfish, while many recently duplicated genes are expressed in an organ-specific fashion (**Figure S8**). Taken together these results imply that, cyclostomes show more limited subfunctionalization or specialisation of expression patterns compared with gnathostomes.

### Gene novelties and losses underlying hagfish character evolution

Hagfishes appear to lack a number of organ systems that are shared across vertebrates, with reduced eyes, limited ossification (no real vertebrae) and sensory organs (no ampullary organs)(Dong and Allison, 2021; Janvier, 2007; Zhang et al., 2009). By using gene family reconstruction incorporating the gene models of the hagfish *E. burgeri* (Yamaguchi et al., 2020), as well as assigning PANTHER classification, we established that hagfish underwent the most extensive loss of genes in any deuterostome lineage with 1,415 missing gene families of which 892 originated in deuterostomes or earlier (**Figure 4c, Figure S7d**). Interestingly, we also estimated rates of paralogue losses in our reconciled trees and while hagfish experienced more severe paralogue losses after WGDs than lampreys, they remain comparable with other gnathostome lineages such as chondrichthyans or teleosts (**Figure S7c**). Therefore, hagfishes stand out as having lost all members of entire gene families, which could be associated with functional losses related to their body plan simplification.

Functional enrichment analysis conducted using lamprey to account for the shared WGD history revealed that some of these absent genes are involved in the development and function of missing hagfish traits (**Figure 4e**). For instance, gamma-crystallins that make up the lenses of vertebrate eyes are absent in hagfish (while independently expanded in lamprey) as is the EYS gene (Eyes Shut homologue) involve in retinitis pigmentosa disease, RBP3 (retinol binding protein 3) expressed in photoreceptor (**Table S9**) (Dong and Allison, 2021).

Hagfish show loss of key genes involved in bone development and hormonal control in other vertebrates. In particular, hagfish lost two members of the RANK/Osteoprotegerin pathway controlling osteoclast proliferation in gnathostomes (Theill et al., 2002) as well as the genes encoding the Parathyroid Hormones (PTH and PTLH) involved in the regulation of calcium metabolism (their receptor is still present) (Poole and Reeve, 2005). These genes are still present in lampreys and were therefore lost in the hagfish lineage, which could be associated with the limited ossification of the hagfish skeleton. Similarly, the hagfish also appears to have a reduced set of genes involved in hormonal control relative to lamprey, affecting both ligand processing (steroid metabolism) and receptors including the absence of the Vitamin D or melatonin receptor. The role of glucagon on glucose metabolism also seems to have been altered with for example the absence of HNF3 (Gauthier et al., 2002).

Hagfish also possess specific adaptations and novelties of which we attempted to uncover some of the genetic bases. Examination of expanded gene families revealed several gene families that are uniquely expanded in hagfish, notably including an extensive expansion (52 copies) of PIF1 helicase homologs, which are involved in DNA repair (Jimeno et al., 2018) (**Table S10)**. Another remarkable characteristic of the hagfish is its prodigious ability to secrete slime, a highly viscous mucus that plays a role in evading predators. We examined genes that are specifically expressed in the slime gland (**Figure S8**) and found two clusters of genes related to intermediate filaments (alpha-keratin) that represent the most highly expressed transcripts in the slime glands (Fudge et al., 2009). Interestingly, one of these clusters contains a gene that is mainly expressed in the skin but not the slime gland, suggesting that the keratin threads of the slime could have originated as elements of the skin, as recently suggested (Zeng et al., 2023) (**Figure 4f**). While slime has been hypothesised to also possess a mucin-type component (Fudge et al., 2005), the most highly expressed glycoproteins are instead related to Von Willebrand factors of type A and D and no mucin-type domains were detected (**Supp. file 1**), suggesting that Von Willebrand factor homologs could potentially serve as an alternative glycoprotein to classical mucins in the context of hagfish slime. These observations suggest that despite their lost characters, hagfish display a number of novelty and adaptations underpinned by their rearranged gene repertoire.

### Programmed DNA (and gene) elimination in hagfish

Hagfish are known to possess distinct karyotypes in their germline vs somatic cells, presumably as a result of the loss of germline-specific regions via embryonically programmed genome rearrangement (as has been described in lamprey embryos) (Bryant et al., 2016; Kohno et al., 1986, 1998; Nakai and Kohno, 1987; Smith et al., 2018, 2012, 2009). Previous studies have identified a large number of satellite repeat elements that are highly enriched on the germline-specific chromosomes/regions, and indicate such elements represent a large fraction of eliminated chromatin across hagfish species (Goto et al., 1998; Kojima et al., 2010; Kubota et al., 1993; Nabeyama et al., 2000). Based on karyotypic and flow cytometry, it was estimated that ∼1.3 Gb was lost from the ∼3.3 Gb *E. atami* germline genome. As yet, no genes have been identified on the eliminated chromosomes of any hagfish species, however,, analyses from lampreys suggest that programmed DNA elimination acts to repress the somatic expression of genes related to germline/pluripotency functions, which may partially resolve fundamental genetic conflicts between germline and soma (Bryant et al., 2016; Smith et al., 2018, 2012, 2009).

To identify regions of the genome that are eliminated by programmed DNA loss, we generated Illumina sequence reads (150 bp paired-end reads) from germline and somatic DNA from a single individual: 117 Gb from testes (∼33X coverage) and 114 Gb from blood (∼54X coverage). Analysis of these reads identified 81 Mb of the assembly enriched in germline relative to soma. These candidate germline-specific intervals contained 1,654 genes, 226 of which had identifiable human homologues (121 non-redundant human genes) (**Table S11**). PCR validation of 46 predicted germline-specific intervals confirmed that 44 could be amplified from testes DNA, but not blood DNA (95.7% validation rate). In both zebra finch and lamprey, it has been observed that somatic gene duplicates are continuously captured by germline-restricted chromosomes, with multiple young somatic paralogs being present on ancient germline-specific chromosomes(Kinsella et al., 2019; Timoshevskaya et al., 2023), therefore it is possible that some PCR markers may not be fully diagnostic with respect to germline specificity.

The nature of germline-specific genes sheds light on the likely function of programmed DNA loss and highlights both shared and divergent aspects of this phenomenon in lampreys vs. hagfish. Ontology enrichment analyses identify several functions that are enriched among germline-specific genes, including functions related to cell cycle, cell motility and chromatin/DNA repair (**Figure 5, Table S13**,**14**). Enrichment of these functions is generally similar to lamprey and lends further support to the idea that eliminated genes perform specific functions within the germline(Smith et al., 2018).

**Figure 5.**
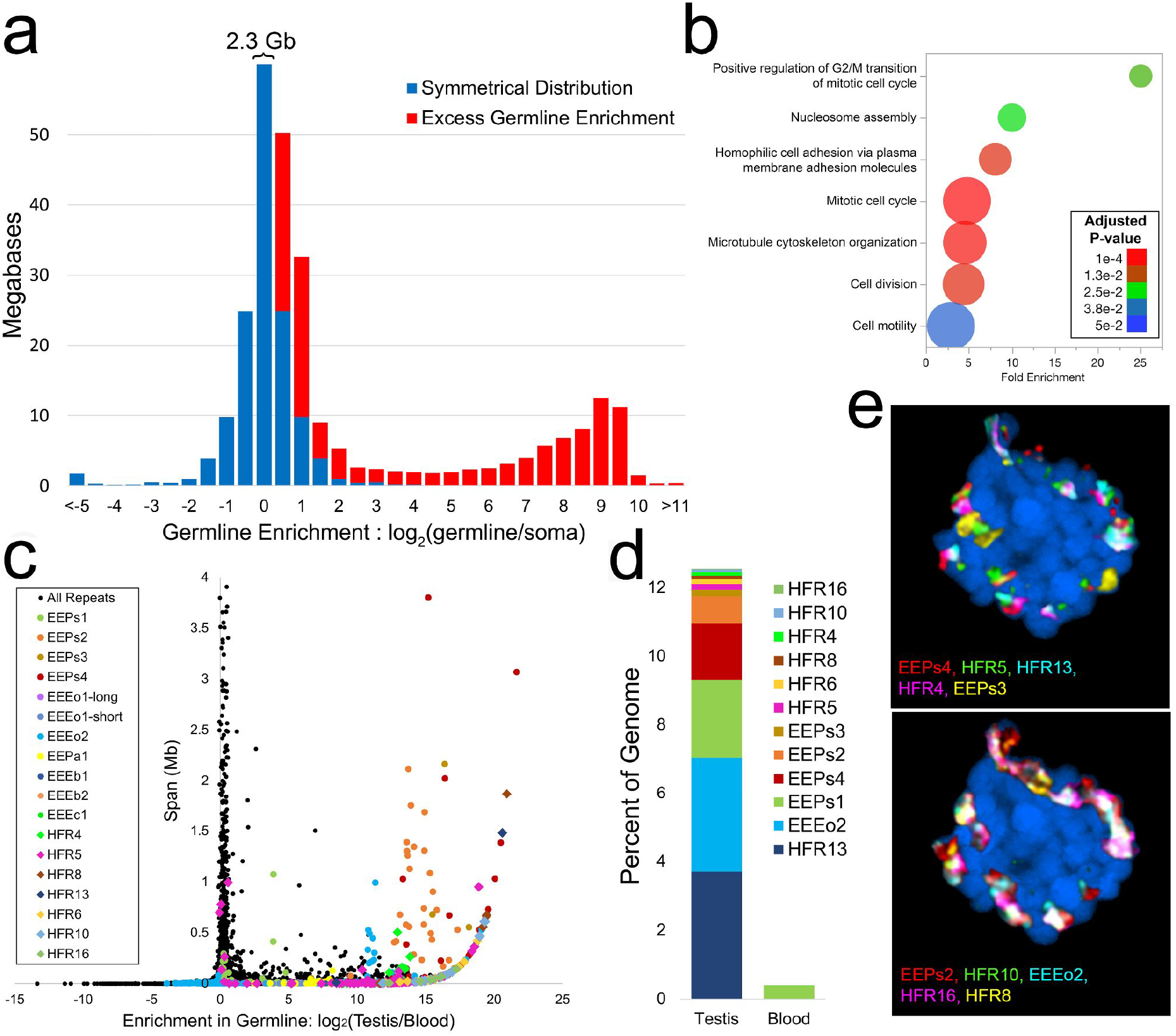
Germline-specific/enriched sequences and genes in hagfish. **a**, Comparison between sequence depth of germline tissue (testes) vs. somatic tissue (blood) identifies a large number of genomic intervals with evidence for strong enrichment in germline. **b**, Genes encoded within germline-specific regions are enriched for several ontologies related to regulation of cell cycle and cell motility (Panther Biological Processes: most specific subclass shown). **c**, Degree of germline enrichment and estimated span of all predicted repetitive elements, focusing on elements with a cumulative span of <4Mb (per family member). Previously identified elements (Kojima et al., 2010; Kubota et al., 1993) are highlighted by coloured circles and newly identified high-copy elements are highlighted by coloured diamonds. Additional higher copy repeats are visible in **Figure S9. d**, Estimated cumulative span of the eight most highly abundant repeats shown as percentage of the genome covered. Colouring scheme is the same as in panels **a** and **b. e**, FISH hybridization of high copy germline-specific repeats to a testes metaphase plate showing their spatial clustering (blue counterstaining is NucBlue: Hoechst 33342; individual pairs of probes are shown in **Figure S11**).

Despite the general functional similarity of eliminated genes in hagfish and lamprey, few orthologous genes were found to be eliminated in both genomes. In total 7 of 121 nonredundant hagfish homologs were also eliminated in sea lamprey (CDH1/2/4, GJC1, MSH4, NCAM1, SEMA4B/C, WNT5/7A/B and YTHDC2, **Figure S9**). Closer examination of gene trees indicates that three of these (orthologs of MSH4, WNT7A and YTHDC2) share a last common ancestor that traces to a single lineage following the basal vertebrate divergence/duplication events. It is possible that this small set of genes may reflect the vestiges of shared germline-specific sequences that were eliminated early in the cyclostome lineage or, alternatively, these genes may have been independently recruited to the germline-specific fraction. Notably, lamprey and hagfish branches for these three genes are generally longer than the branches for their gnathostome homologs. As such, it is also possible that some germline-specific genes have diverged to the point that the correct homologs are not readily identified via automated orthology and tree construction pipelines with the current taxonomic sampling. It seems likely that characterisation of germline-specific genes in other lampreys, hagfish and gnathostome genomes will better resolve the evolutionary origins of programmed DNA loss in vertebrate lineages.

Another contrast between hagfish and lamprey eliminated genes is revealed by analyses of binding sites near their human and mouse homologs in published chromatin immuno-precipitation data (Xie et al., 2021). While these analyses rely on data collected from divergent gnathostome species, we only expect to observe enrichment for ancestrally-conserved functions, and in practice we observe that enriched pathways are generally related to regulation of early embryonic transcription. Homologs of lamprey eliminated genes are highly enriched for targets of Polycomb Repressive Complex (PRC) regulation in embryonic stem cells(Smith et al., 2018), whereas homologs of eliminated hagfish genes do not show evidence for enrichment of PRC targets. Rather, eliminated genes are highly enriched for targets of SMAD2/3 nuclear pathway, nuclear beta-catenin signalling, and neural crest differentiation (**Table S15**). The differences in the collection of genes and specific gene functions that have been relegated to the germline-specific fraction in hagfish vs lamprey may not be particularly surprising given the ∼500 million years that have passed since hagfish and lamprey last shared a common ancestor, and the dramatic differences in ecological (benthic scavengers *vs*. anadromous filter feeders/ectoparasites), developmental, and reproductive (iteroparous *vs*. semelparous) biology.

In both hagfish and lamprey, the eliminated chromosomes contain large numbers of highly repetitive satellite sequences. These sequences are often not fully incorporated into genome assemblies and result in increased fragmentation of chromosomal assemblies for germline-specific regions (Timoshevskaya et al., 2023). To more fully characterise arrays of repetitive elements in hagfish we assembled satellite and other repetitive sequences directly from the reads (their component k-mers) and merged these with a library of repeats that was compiled from the assembly. Aligning germline and somatic readsets to this collection of repetitive elements allowed us to calculate the relative copy number of repeats in germline vs soma and estimate the combined span of each repeat within the genome (**Figure S9D**). Many of these individual repeats could be clustered into repeat families consisting of highly similar (>%80 nucleic acid identity) elements. These families included several satellite elements that were previously identified as enriched within the germline (Goto et al., 1998; Kojima et al., 2010; Kubota et al., 1993; Nabeyama et al., 2000), and other elements that had not been previously identified. PCR validation confirms the presence of both new and previously-predicted satellite repeats as well as the degree to which they are specific to germline, and FISH hybridization to testes and somatic cells further verifies that these are highly enriched in germline, linked in interphase nuclei, and distributed across several distinct germline-specific regions (chromosomes) (**Figure S10D** and **S11**). Among our newly identified set of germline-specific repeats was the most abundant germline element yet to be identified in *E. atami* (HFR13) a 67 bp tandem repeat that is estimated to span >128 Mb, or ∼3.7% of the germline genome (>9% of the germline-specific fraction). In total, the 12 highly abundant and validated repeats reported above are estimated to span more than 30% of the germline-specific fraction (**Figure 5**).

## Conclusion

With the description of the hagfish genome and confirmation of cyclostome monophyly, it is now finally possible to infer the full sequence of genomic events that shaped early vertebrate evolution. We found that despite their distinct chromosomal architectures, hagfish and lamprey share the same genome duplication history, with a shared hexaploidisation event in the cyclostome lineage (Nakatani et al., 2021; Simakov et al., 2020; Smith et al., 2018). The multitude of lamprey chromosomes, therefore, reflects the ancestral cyclostome condition, in the same way, that the large chromosome number of chondrichthyans reflects the ancestral state within gnathostomes (Marlétaz et al., 2023; Nakatani et al., 2021). The hagfish lineage experienced extended fusions and rearrangements from these ancestral duplicated chromosomes (Figure 2). Our molecular dating analyses suggest that the emergence of vertebrate-specific characters(Shimeld and Holland, 2000) and the associated subfunctionalization of paralogous genes took place in a relatively short period between the diploidization following 1R_V_ and the cyclostome-gnathostome divergence (**Figure 3**).

The preferential involvement of 1R-paralogues with functional terms classically associated with vertebrate emergence corroborates this view (**Figure 4a**). For instance, we detected a significant enrichment of 1R-paralogues with terms associated with neural and embryonic development and we noticed that, among adult tissues, such 1R-paralogues are preferentially expressed in the nervous system (**Figure S8**). For instance, many of the genes involved in the specification of the neural crest seem to have emerged and acquired their function early after the 1R_V_ duplication in vertebrate evolution (Martik and Bronner, 2021), but also genes involved in hormonal control(Kuraku et al., 2023) (Fig. 4a). We cannot rule out, however, the possibility that delayed rediploidisation, as observed in teleosts (Parey et al., 2022), could have eroded the phylogenetic signal associated with WGD events and distorted molecular dating analyses. Nevertheless, an early acquisition of gene specialisation associated with vertebrate characters is consistent with the involvement of these novel genes in the vast radiation of ostracoderms (Janvier, 2015).

The cyclostome lineage emerged from early vertebrate diversity and experienced an additional lineage-specific hexaploidisation event, but there appears to have been more gene loss and less subfunctionalization in cyclostomes than in gnathostomes (Marlétaz et al., 2018; Nakatani et al., 2021). Nevertheless, some hagfish-specific novelties are evident in the genome, particularly as they relate to the evolution of slime and DNA repair. The evolution of embryonically programmed DNA loss led to the evolution of large numbers of novel lineage-specific coding genes that are only present only in the genomes of germ cells in lampreys (Smith et al., 2018) and hagfishes (**Figure 5**). We speculate that the presence of these genes might have introduced constraints at the level of genome regulatory architecture related to the distinct modalities of paralogue evolution in cyclostomes vs. gnathostomes. Notably, hagfish are characterised by accelerated evolution relative to lampreys. At the karyotype level, they sustained extensive chromosomal rearrangements and fusions of ancestral chromosomal elements (**Figure 2**), while at the morphological level, they suffered prominent character loss (eyes, ossification, sensory organs), which is strikingly associated with associated gene losses. Our findings suggest that vertebrates are not immune to evolutionary trends that were not so long ago thought to be restricted to some of the most derived lineages, for instance, appendicularians (Ferrández-Roldán et al., 2021), and reaffirm the importance of gene loss as an evolutionary force (Albalat and Cañestro, 2016).

## Supplementary Figures (S1-S11) Tables (S1-S16)

### Supplementary files

**Supplementary File 1**. Table gathering essential information for each gene model in *E. atami* including location, protein domains, gene family, gene expression cluster [Eptata_genes_filt.xlsx]

**Supplementary File 3**. Phylogenetic trees inferred for paralogons in each CLG assuming the C20+R model [paralogons_t6_c20.pdf]

**Supplementary File 3**. Phylogenetic tree of cadherin-related genes showing the tandem duplication patterns in gnathostomes [OG_4678_Cadh.tre.pdf]

**Supplementary File 4**. Functional enrichment for sets of paralogues showing distinct retention pattern after genome duplications in vertebrates (Figure 4b). [Vert2R_Go_enrich_wg_gds.txt].

**Supplementary File 5**. Functional enrichment in lamprey for gene families lost in hagfish (data for Figure 4e). [Loss_Lamprey_GO_gn_dsc.tsv]

**Supplementary File 6**. Synteny-based paralogue classification for reconstructed gene families [Vert_Evt_OGrrA.txt]

## Author contributions

D.S.R, O.S., F.M., J.S., and S.B. conceived the study, which was led by F.M., J.S., and D.S.R. F.M. and O.S. sequenced, assembled, and annotated the genome. N.T., V.T. and J.S. performed the DNA elimination analysis. F.M., O.S. and D.S.R. performed synteny analyses. FM. completed phylogenetic and paralogon analysis. E.P. contributed duplication phylogenetic modelling and Hox cluster analysis. D.G. analysed transcriptomic data. M.S. provided biological samples and figure material. D.S.R., F.M., and J.S wrote the paper with input from N.T., V.T. and E.P. All authors read and approved the manuscript.

## Acknowledgement

We thank B. Venkatesh and S. Kuraku for early discussions, K. Kubokawa for discussion of hagfish sampling, N. Segi, T. Suzuki, B. Muramatsu, and H. Dohra for technical support; and H. Hasegawa and K. Hasegawa for hagfish supply. Work at the OIST Molecular Genetics Unit (D.S.R., O.S., D.G. and F.M.) was supported by OIST internal funds. F.M. is supported by the Royal Society Fellowship URF\R1\191161 and the BBSRC grant BB/V01109X/1. D.S.R is a Chen-Zuckerberg BioHub Investigator and is supported by the Marthella Foskett Brown Family Chair of Biological Sciences at U.C. Berkeley. J.S. is supported by grants from the National Institutes of Health (NIH) (R35GM130349) and National Science Foundation (NSF) (MCB1818012). M.S. was in part supported by the Field Science Center and Research Institute of Green Science and Technology, Shizuoka University. E.P. is supported by a Newton International Fellowship from the Royal Society (NIF\R1\222125). We thank the OIST Sequencing Section for DNA and RNA sequencing and acknowledge the support of OIST supercomputing and computational support from the University of Kentucky High-Performance Computing complex

## Methods

### Genome sequencing and assembly

DNA was extracted from a testis from a male *Eptatretus* (formerly *Paramyxine*) *atami* individual and extracted using protein K digestion and phenol:chloroform extraction (Green and Sambrook, 2012). Animals were anaesthetized using Tricain (MS222, Sigma) prior to sacrifice and dissection. Paired-end and mate-pairs illumina libraries were generated using Illumina Truseq and Nextera Mate-pair kits and sequenced on HiSeq2000 and HiSeq2500 instruments (**Table S1**). The illumina dataset was assembled using meraculous (v2.2.2.5) with a k-mer of 71 and ‘diploid mode’ set to ‘1’ to attempt the merging haplotypes (Chapman et al., 2011)and subsequently scaffolded using mate-pairs information (Table S2). PacBio long-reads data at ∼35x coverage were generated on a PacBio RSII instrument (Table S2) and incorporated using PBJelly (v15.8.24) (English et al., 2012). PBJelly aligns the PacBio reads to the assembly using the Blasr aligner and collects reads surrounding and spanning gaps. Sequences assembled from these spanning reads are used to fill gaps and extend scaffolds. We used the parameters ‘-minMatch 8 -sdpTupleSize 8 -minPctIdentity 75 -bestn 1 - nCandidates 10 -maxScore -500’ for Blasr alignment.

The gap-filled assembly was further scaffolded using proximity ligation information. We used both Chicago libraries relying on syntenic reconstructed chromatin and HiC libraries capturing the native chromatin contacts and scaffolding was performed using the HiRise package (Putnam et al., 2016). Hagfish liver was crosslinked in 1% PFA, and chromatin subsequently extracted, immobilised on SPRI beads, washed and digested with DpnII (Meyer and Kircher, 2010). After end-labelling, proximity ligation was carried out using T4 DNA ligase and cross-linking reversed using Proteinase K, removed from the beads and the DNA fragments were purified again on SPRI beads. Sequencing library was constructed using the NEB Ultra library preparation kit (New England Biolabs, Ipswitch).

The genome polymorphism was estimated to be 0.9%. The final BUSCO score (Metazoa) is C:90.0%[S:89.8%,D:0.2%], F:4.0%, M:6.0%, n:954. The size of the hagfish genome was estimated by counting 21-mers with Meryl (v1.1) (Miller et al., 2008). Using a fitting 4-peak model as implemented in Genomescope2, the estimated size is 2.02Gb and 3.28Gb using sequencing data from blood and testis DNA, respecting (**Figure S1b**) (Vurture et al., 2017).

### Transcriptome and genome annotation

We generated RNA-seq data for 13 organs with 26M reads on average (**Table S8**). We aligned the reads to the genome using STAR (v2.5.2b) with an average 78.7% uniquely mapping reads (Dobin et al., 2013). These alignments were employed to assemble transcriptomes for each organ using Stringtie (v1.3.3b) and subsequently merged together using Taco (Niknafs et al., 2017). In parallel, a *de novo* assembly of the bulk RNA-seq data was performed using Trinity both in reference-free and genome-guided mode (Grabherr et al., 2011).

We also sequenced full-length cDNA from Brain RNA on 8 cells of Pacbio RSII. Following the Iso-Seq protocol, circular consensus (CCS) of subreads were calculated and validated as full-length based on the presence of SMART adaptors at both extremities. Full-length transcripts were clustered and polished using all ccs reads with quiver (v2.0.0), yielding 23,343 high-quality transcripts.

Assembled transcripts from *de novo* and genome-guided Trinity and high-quality isoseq transcripts were aligned to the genome using GMAP (v. 2018-03-25). Mikado (v1.2.1) was used to generate a high-quality reference transcriptome leveraging (i) the aligned trinity *denovo* and genome-based transcriptomes, (ii) the isoseq transcripts, (iii) the stringtie transcriptomes merged with Taco and a set of curated splice-junctions generated from RNA-seq alignments using Portcullis (v1.0.2). Putative fusion transcripts were detected by Blast comparison against Swissprot and ORFs were annotated using Trans-decoder (Haas et al., 2008). Transcripts derived from the reference transcriptome were selected to train the Augustus *de novo* gene prediction tool (Stanke et al., 2006). Intron positions and exon positions were converted into hints for Augustus gene prediction.

Finally, we constructed a database of repetitive elements using RepeatModeler (v1.0.11) and employed it for masking repetitive sequences with RepeatMasker (v4.0.7). Gene models with half or more of their exons showing 50% overlap with repeats were discarded, yielding 46,822 filtered gene models. Alternative transcripts and UTRs were subsequently incorporated using the PASA pipeline (Haas et al., 2008). These gene models contain a total number of 4915 distinct PFAM domains.

### Phylogenomics and molecular dating

We inferred a set of 1562 single-copy orthologues suitable for phylogenetic reconstruction by applying the OMA tool (v2.4.1) (Altenhoff et al., 2019) to a subset of deuterostome proteomes including lamprey and the newly generated hagfish gene models (Table S6). Selected transcriptomes were assembled using Trinity and translated using transdecoder (v5.5.0) (Haas et al., 2008). We built HMM profiles using Hmmer (v3.1b2) for each orthologue family and extracted orthologues for phylogenetic reconstruction using the same approach as in (Marlétaz et al., 2019). Subsequent sequences were aligned using Msaprobs (Liu et al., 2010), mistranslated stretches filtered out using HmmCleaner (Di Franco et al., 2019) and diverging regions intractable for phylogenetic analysis removed using BMGE (-g 0.9) (Criscuolo and Gribaldo, 2010). Phylogenetic trees were reconstructed for each alignment using IQ-TREE (v2.1.1) with a LGX+R model (Minh et al., 2020). For computationally intensive analyses, such as site-heterogenous reconstruction with CAT+GTR, we selected the 20% orthologues with the lowest saturation. Molecular dating analysis was conducted using Phylobayes (v4.1e) (Rodrigue and Lartillot, 2014) using the CAT+GTR+G4 model and the CIR relaxed clock (with soft-bound) assuming fossil calibrations (**Table S4**) (Irisarri et al., 2017; Kuraku et al., 2009b; Miyashita et al., 2019).

### Synteny reconstruction

Pairs of orthologous genes were obtained by mutual-best-hit after reciprocal proteome comparison using MMSeqs2 (r12-113e3) and were used to create a system of joint coordinates to plot orthologue position in two species. Fisher’s exact test was used to determine mutual enrichment of orthologues between chromosomes, and only significant enrichments were incorporated in binned orthologous content representations (Fig. 2a). Plots connecting orthologues in multiple species (Fig. 2b) were generated using Rideogram (v0.2.2).

### Gene family analyses and phylogenetic analyses of paralogons

We reconstructed gene families using Broccoli (Derelle et al., 2020) for a set of genomes from deuterostome species (Table S6). For gene families that included at least 6 genes and 3 species but less than 450 sequences in total, we applied Generax to infer the losses and duplications that affected a given gene family (Morel et al., 2020). To do that, we generated individual alignments using MAFFT (v7.305) (Katoh and Standley, 2013), filtered them using BMGE and reconstructed a tree using IQ-TREE and a LG+R model (Minh et al., 2020). These curated alignments and trees were used as input for Generax (v1.2.2) assuming a D+L (duplication plus loss model). Reconciled trees in the RecPhyloXML format were parsed to estimate the duplications and lineage-specific losses at each node of the species tree as seen in Figure S6 (Duchemin et al., 2018). Reconciled trees were split if they showed a duplication at the ‘deuterostomia’ node indicative of a deep paralogy relationship.

For each gene family, we first assigned the CLG by considering the location of amphioxus and sea urchin genes and the corresponding CLG-to-chromosome assignment, and then evaluated the occurrence of the paralogues derived from the 1R and 2R_jv_ in gnathostomes based on the vertebrate classification previously established(Simakov et al., 2020) and revised in this study (**Table S6**). Selected species for gene families including derivatives of the 1R paralogons and at least 3 out of 4 possible paralogons for gnathostomes (α1, α2, β1, β1) were collected (**Table S7**). These genes were concatenated for each CLG based on their paralogon identity in gnathostomes, and the chromosomal identity of the CLG derivatives in cyclostomes. Two datasets were generated, a ‘strict’ one where at least 3 distinct gnathostome paralogons were required for each retained gene family and a ‘relaxed’ one where only two or more gnathostome paralogons were required (**Table S6**). A similar approach was used to classify individual genes depending on the duplication events from which they derive **(Supp. file 3**). We collected Gene ontology terms and functional classification information by applying eggnog (Huerta-Cepas et al., 2019) on the proteome of our interest species and term enrichment analysis conducted using the TopGO package (v2.50.0).

### Tests of WGD hypotheses on the vertebrate phylogeny

We used the WHALE software v2.1.0 (Zwaenepoel and Van de Peer, 2020) to rigorously test WGD hypotheses on a reduced vertebrate species tree (Fig. S4). We leveraged a total of 8,931 gene families in this analysis, selected to contain at least one gene copy in each clade from the root, in compliance with the assumption of WHALE that genes were acquired in a common ancestor of all included species. We further filtered large families to reduce the computational burden. For each of the 8,931 retained families, we built a multiple sequence alignment based on the amino acid sequences with MAFFT v7.508 (Katoh and Standley, 2013) and reconstructed 1000 bootstrap trees with IQ-TREE 2.2.0.3 (Minh et al., 2020) under the LG+G model. We summarised clade conditional distribution (CCD) from bootstrapped trees using the ALEobserve tool from the ALE software (Szöllõsi et al., 2013). We ran WHALE on the dated species trees and CCD data to test 5 WGD hypotheses on the vertebrates species tree: the 1R in the vertebrate ancestor, the 2R in the gnathostome ancestor, a cyclostomes-specific duplication, a hagfish-specific duplication and a lamprey-specific duplication. We used the variable rate DLWGD WHALE model, which models independent duplication and loss rates across branches. We assumed a normal distribution N(log(0.15), 2) on the mean log-scaled duplication and loss rate, an exponential distribution (mean=0.1) prior on its variance, a Beta (3, 1) hyper prior on the η parameter (distribution of the number of genes at the root), and uniform priors on the retention parameters (q parameter) for all WGDs. We obtained significant Bayes factors (BF_Null_vs_WGD_ < 10^−3^) in support of large-scale duplication (q parameter ≠ 0) for the 1R, 2R and cyclostomes-specific events. These results were reproduced using the simpler constant rate DLWGD model.

### Phylogenetic tree based on hox clusters concatenation

We investigated the phylogenetic relationships between hox clusters and bystander genes in 8 genomes: amphioxus, sea lamprey, hagfish, human, mouse, chicken and spotted gar. We identified hox and bystander genes in three steps: (i) starting from human gene names, we searched for orthologs in the other species using our set reconciled gene trees (generax trees), (ii) we used NCBI blastp (Johnson et al., 2008) to confirm identified hox genes and further search for hox genes missed by the gene trees approach, (iii) we used miniprot 0.5-r179 (Li, 2023) with the sets of human and *E. burgeri* hox proteins to search for hox genes missing from genome annotations of other species. We next aligned each gene family using their amino-acid sequence with MAFFT v7.508 (Katoh and Standley, 2013) and concatenated alignment from each cluster. Finally, we used the concatenation matrix to build a phylogenetic tree with RAxML-NG v. 1.1 (Kozlov et al., 2019) using the LG+G4+F model, 10 different starting parsimony trees and 100 bootstrap replicates.

### Comparative transcriptomics

RNA-seq reads for hagfish (this study), the lamprey *Lampetra japonica* (PRJNA354821, PRJNA349779, PRJNA312435), the gar *Lepisosteus oculatus* (PRJNA255881) and the cephalochordate amphioxus (PRJNA416977) were aligned with STAR (v2.5.2b) (Dobin et al., 2013), counts for annotated genes obtained using featureCount from the subreads package (v1.6.3)(Liao et al., 2014). Counts were converted to FPKM in the R package for subsequent analyses: WGCNA (v1.7.0) was used to cluster gene expression in the full organ set: after filtering out genes with limited variance and coverage, the ‘softpower’ parameter was estimated to 20, and clustering was run with a ‘signed’ network type (Langfelder and Horvath, 2008). The gene expression specificity index (or *τ*) was calculated as described in (Yanai et al., 2005) on sets of organs (brain/neural tube, gills, heart, intestine, kidney, liver/hepatic tissues, ovaries/female gonad, skin/epidermis and muscle). For comparative analyses, gene families with paralogues derived from the vertebrate WGD were selected based on their duplication history, and the gene expression specificity index was compared across species for the same gene families (**Figure S7**). We also compared gain and losses of expression domains for a given gene family by binarising gene expression across a reduced set of 6 organs (brain, gills, intestine, liver, muscle, ovary) and counting expression patterns gains of lost between genes belonging to a given gene including paralogues and outgroup. The number of gain and loss events is then plotted as a distribution centred around zero (**Figure 4d**).

### Detection of Germline-Enriched/Specific Regions

DNA was extracted from testes and blood via phenol-chloroform extraction(Green and Sambrook, 2012). In order to enrich for germ cells, testes tissue was ground gently with a plastic pestle in a 1.5 ml microfuge tube and residual connective tissues were discarded prior to Proteinase K digestion. Outsourced library prep and Illumina sequencing (HiSeq2500 V4, 150 bp paired-end reads) was performed by Hudson Alpha Genome Services Laboratory.

Sequence data were aligned to the *E. atami* genome assembly using BWA-mem (version 0.7.5a-r416)(Li and Durbin, 2009) with option -a and filtered by samtools view(Li and Durbin, 2009) with option -F2308. Only primary alignments with mapping quality 5 and higher were retained for further analysis. The resulting files were processed using DifCover (Version 4)(Smith et al., 2018) to calculate the degree of germline enrichment across all discontiguous 500 bases intervals of low-copy sequence using modal coverages for sperm and blood of 32X and 54X respectively, low coverage masking of regions with read depth <1/3X in both samples and high coverage masking of sequences with read depth >3X modal coverage in both samples. To identify germline-specific genes that are present at higher copy number, we ran DifCover using low coverage masking with read depth <10X in both samples and high coverage masking of sequences with read depth >30X modal coverage.

### PCR validation of germline-enriched loci

Primers were designed using a coverage-masked version of the *E. atami* genome. using Primer3(Koressaar and Remm, 2007) (version 4.1.0). PCR validation reactions amplification were performed using GoTaq® DNA polymerase (Promega, 1.2 units/50 μl reaction), Colorless GoTaq® Reaction Buffer, 1 μg of genomic DNA template and 100 ng of oligonucleotide primer. PCR cycling conditions included a 3 minute initial denaturation step at 95 °C, 34 cycles of a three-step thermal cycling consisting of a 30 second denaturation at 95 °C, a 30 second primer annealing step at 55-65 °C (**Table S12**), and a 30 second extension step at 72 °C. A final extension at 72 °C was performed on all reactions to ensure production of full length amplicons. Amplification was assessed by agarose gel electrophoresis. Eight primer pairs with ambiguous signal in the first rounds of PCR were redesigned and retested (**Table S12**). We note that some PCR markers may not be fully diagnostic with respect to germline specificity, since, as observed in zebra finch, somatic gene duplicates have been continuously captured by the germline restricted chromosome since its presumptive origin within the ancestral songbird lineage(Kinsella et al., 2019)).

### Computational prediction of germline-enriched and highly abundant somatic repeats

Abundant *k*-mers (*k* = 31) were identified from testes and blood DNAseq sequence data using Jellyfish version 2.2.4(Marçais and Kingsford, 2011). Minimal copy-number thresholds for defining abundant *k*-mers were set at 3X the modal copy number: 72 for testes and 120 for blood. Abundant *k*-mers were extracted and assembled into a set of *de novo* assembled repetitive sequences using Velvet version 1.2.10(Zerbino and Birney, 2008) with a hash length of 29. These sequences were aligned (blastn with -word_size 17) to repetitive elements generated from *E. atami* genome assembly by RepeatModeler (Smit and Hubley, 2008) and sequences that aligned with <90% identity or under 80% of their length were added to the set of reference-derived repeats to form a union set.

Enrichment analysis was performed by separately aligning paired-end reads from testes and blood to the union set. Primary alignments, identified by samtools view (Li et al., 2009) with option -F2308, were additionally filtered to retain only alignments that either cover >80% of a repeat or have >80% of read bases aligned. Enrichment scores were calculated with DifCover pipeline v.3 (Smith et al., 2018). Stage2 of the pipeline was run with parameters v=10000, l=0, a=b=10, A=B=10_8_. Stage 3 of the pipeline was modified by employing a subroutine from DNAcopy(Seshan and Olshen, n.d.) without “smoothing” the data prior to analysis. From a set of 180,032 intervals generated by DifCover we chose 138 highly abundant and germline-specific sequences with enrichment scores of more than 10 and estimated span size of more than 100 Kb. The estimated genomic span of these repeats was computed as [(testes coverage/modal testes coverage)X(number of bases with read depth coverage > 10)], where modal testes coverage = 32. Clustering of 138 highly abundant and germline-specific sequences was performed using CD-HIT-EST (v4.6, with parameters: -c0.8, -G0, -aS 0.3, -aL 0.3, -sc 1, -g 1, -b 4)(Li and Godzik, 2006) resulting in the formation of 38 clusters that were further merged to 24 by manual curation and cross alignment of sequences from the initial clusters. For characterization of repetitive structures and identification of motifs representative sequences from each cluster were mapped to the assembly (blastn, -word_size 15) and to a collection of published hagfish repeats. We found that four of 24 clusters have sequences that are homologous to the published repeats of *Paramyxine sheni* EEPs2, EEPs3, EEPs4 and *E. okinoseanus* EEEo2(Kojima et al., 2010; Kubota et al., 1993). Primers for these and representatives of 7 other clusters were designed with Primer3 (v. 0.4.0) tool (**Table S16**).

To facilitate FISH visualisation, we also searched for possible candidates for centromeric repeats. Such candidates are expected to be 1) highly abundant in both somatic and germline sequence, and 2) be enriched in a “centromeric” region of every chromosome. From the union set we chose repeats with blood coverage > 10_5_ or span > 1Mb and aligned them to the assembly (blastn - word_size 15, p>75, coverage > 80%). Repeats with more > 200 hits within a 1 Mb window were grouped to three families labelled Soma1-3. Soma2 appeared to be homologous to *P. sheni* repeat EEPs1(Kojima et al., 2010), Soma1 and Soma3 to *E. burgeri* contigs LC047612.1 and LC047003.1. FISH analysis confirmed that as it was predicted *in silico* EEPs1 is highly abundant in both testes and blood DNA of *E. atami*.

To estimate more accurately genomic span of chosen germline enriched and somatic repeats we realigned reads from blood and testes to the sequences of these repeats or to the sequence extended as a tandem repetition of repeat’s motif spanning at least 150 bps (**Table S12**) and applied all described previously steps for filtering and coverage and span estimation.

### *In situ* Hybridization

#### Slide preparation

Snap frozen blood and testes samples were used for slide preparation of somatic and germline cells for validation of the presence and specificity repeats in different cell types. A small amount of blood (about 20 mg) was gently thawed on ice, mixed with 2 ml buffered hypotonic solution (0.4% KCl, 0.01M HEPES, pH=6.8), and incubated for 30 min at room temperature. The cells were prefixed by gentle mixing suspension with several drops of fixative solution (methanol:acetic acid - 3:1). After centrifugation (5000×g for 10 min) supernatant was removed and cells were resuspended and fixed with methanol:acetic acid - 3:1. Three additional fixative solution changes were carried out to ensure cells were fully equilibrated to fixative solution. Fixed cells were stored at -20 °C. One fixative change was made before spreading the cell suspension onto slides. About 20 mL drop was applied on a steamed slide which was immediately placed on a heating block in a humidity chamber at 60 °C for 1-2 min. After air drying, slides were examined with a microscope using a low condenser position to aid in viewing unstained nuclei/metaphases. Slides were aged 1-3 days prior on a warming stage at 37°C prior to hybridization. For germline cells, a piece of testes (30-40 ng) was minced with a razor blade, placed in a homogenizer, and disaggregated in hypotonic solution. Testes cell suspensions were filtered through 40-50 mm cell strainer to remove excess tissue. Subsequent steps of fixation and slide preparation for testes tissue were as described for blood.

#### Probe labellin

Probes for FISH were generated using a modified conventional PCR: the reaction mix with final volume of 25 μL contained 0.1 mM each of unlabeled dATP, dCTP, and dGTP and 0.03 mM of dTTP; 0.5 μL one fluorophore conjugated dUTP (Cyanine 3-dUTP (Enzo), Cyanine 5-dUTP (Enzo) or Fluorescein-12-dUTP (Thermo)), 1× Taq-buffer and 0.625 U GoTaq DNA Polymerase (Promega). Each PCR amplification was performed using 0.5μg of genomic DNA template from testes, 34 PCR cycles and a 30 second extension step to obtain appropriately sized probes for FISH. After cycling reaction 25 μL of PCR mix were combined with 5 μL sheared salmon sperm DNA (1 mg/mL, Thermo), 3 μL 3M Sodium Acetate, pH=5.2, and 80 μL 100% cold ethanol and kept overnight at -20 for probe precipitation. After spinning and supernatant removal, the pellet was dissolved in 25-30 μL of 50% formamide and stored at -20 °C prior to use.

#### Fluorescence in situ hybridization (FISH)

FISH on chromosome preparations was carried out according to a standard protocol for chromosome spreads(Rooney, 2001) with modifications(Timoshevskiy et al., 2012). Prior to hybridization slides were incubated in 2×SSC for 30 min at 37°C, passed through ethanol series (70, 80, 100%), dried and denatured in formamide (70% in 2×SSC) for 2 minutes, prewarmed to 70 °C. After the formamide denaturation, slides were placed immediately in cold (−20 °C) 70% ethanol, further dehydrated in 80 and 100% ethanol, and kept on slide warmer at 37°C until hybridization mix with probe was applied.

Differently labelled hybridization probes were mixed (1 μL of each per slide) with hybridization master mix (60% formamide, 10% dextran sulphate, 1.2×SSC) to a final volume 10 μL. Hybridization mix was denatured at 95 °C for 7 minutes, cooled in ice, prewarmed to 37 °C, applied to the slide, coverslipped and sealed with rubber cement. After overnight incubation in a humidity chamber at 37°C slides were washed in 0.4×SSC, 0.3% NP-40 for 3 minutes at 70°C and in 2×SSC, 0.1% NP-40 for 5 minutes at room temperature. One drop of ProLong™ Glass Antifade Mountant with NucBlue™ Stain was placed in the center of an area to be examined and covered with a coverslip.

#### Microscopy and image analysis

Slides were analyzed with an Olympus-BX63 microscope using filter sets for DAPI, FITC, Cy3, and Cy5. Images were captured using CellSence software (Olympus) and processed with Adobe Photoshop CC 2019 and ImageJ 1.53k (NIH).

