## Supplementary figures for "The hagfish genome and the evolution of vertebrates"

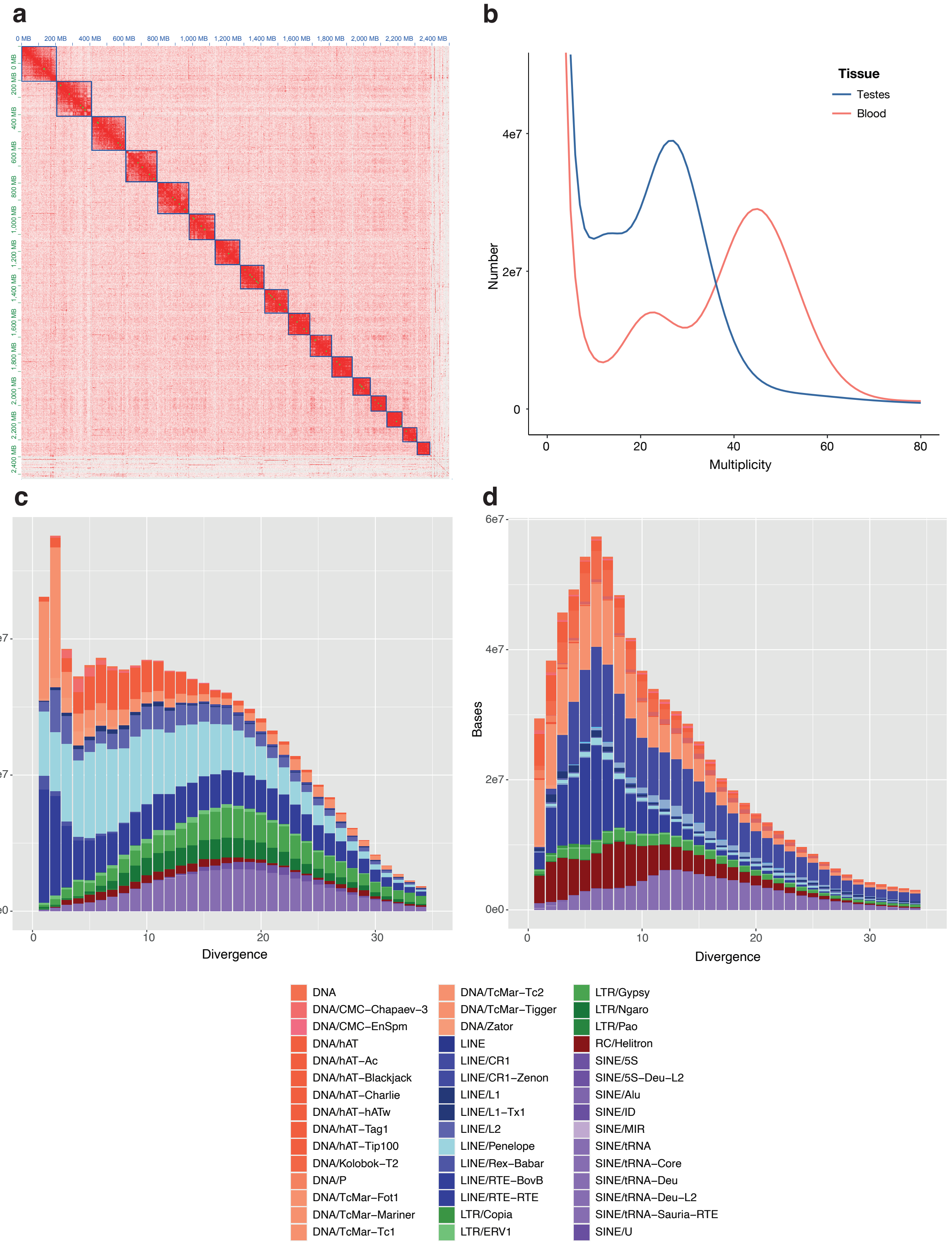


**Figure S1.** Genome content and architecture of E. atami. (**a**) Hi-C contract map highlighting the 17 somatic chromosomes (**b**) 21-mer profile of somatic (blood) and germinal (testes) genome showing estimated genome size of 2.02 and 3.28 Gb, respectively. (**c,d**) Repeat landscape in lamprey (**c**) and hagfish (**d**) showing a markedly different profile.

*Supplementary figures*

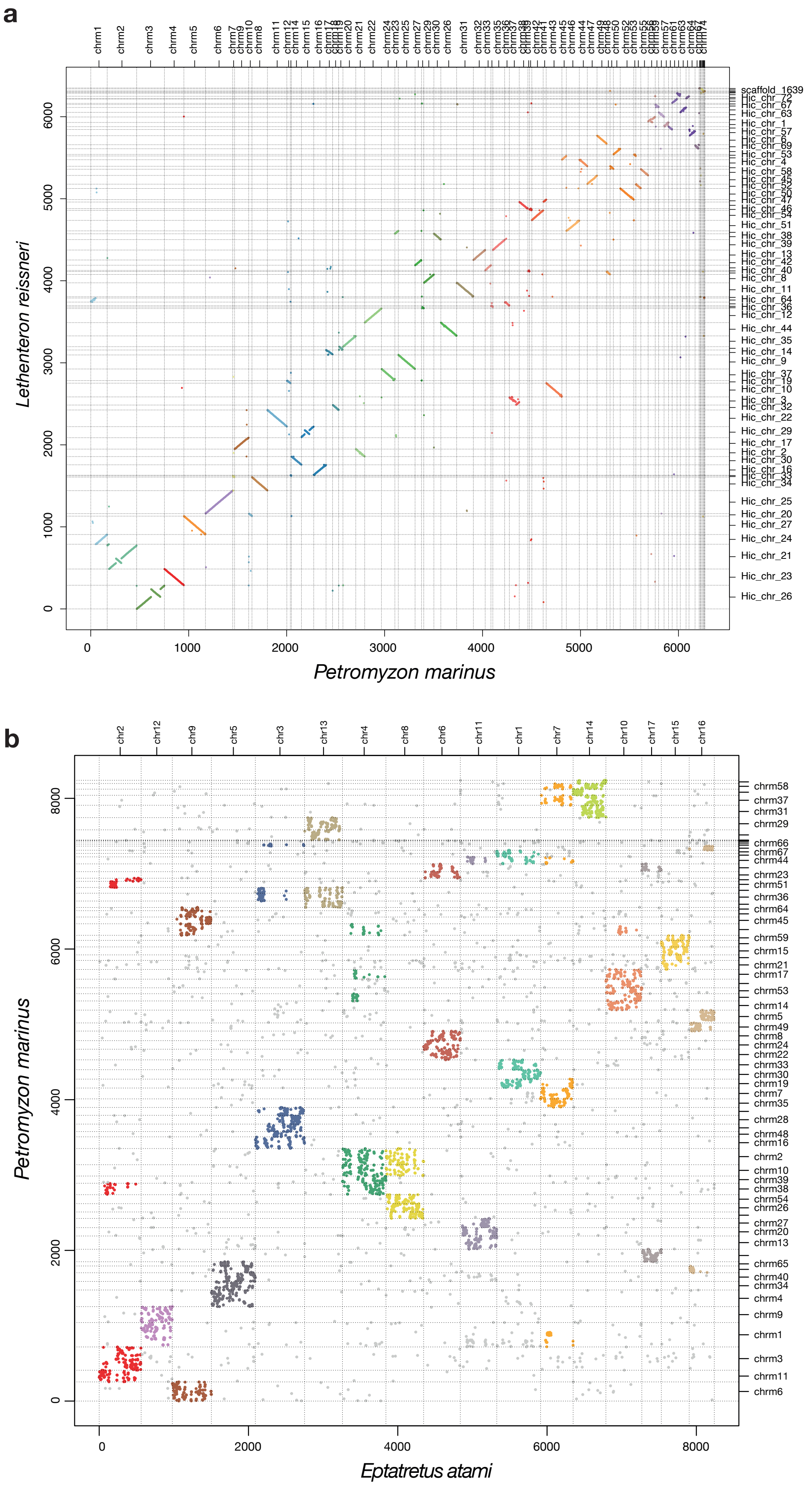


**Figure S2.** Dotplot comparison of chromosomal architecture between (**a**) two lampreys (*L. reissneri* and *P. marinus*) an (**b**) the hagfish *E. atami* and the lamprey *P. marinus*.

*Supplementary figures*

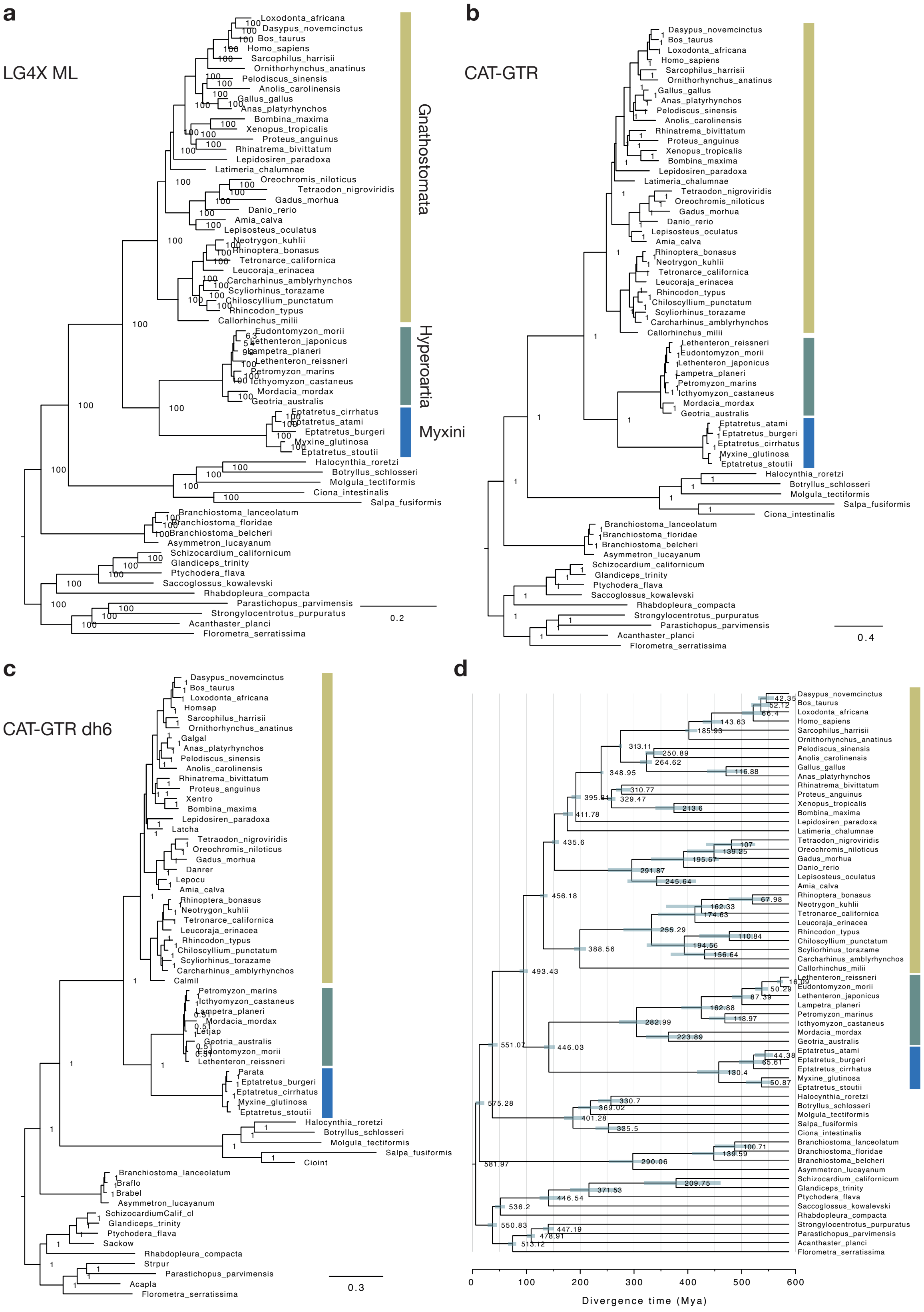


**Figure S3. Phylogenetic reconstruction of deuterostome relationship with a focus on cyclostome positions** (**a**) Tree reconstructed with IQ-TREE assuming LG4X model using a set of 1,467 single-copy orthologues and a partitioned model (**b**) Tree reconstructed using Phylobayes and a CAT+GTR+G4 model using a subset of 176 orthologues showing the lowest saturation (see methods). (**c**) Tree reconstructed using the same set of orthologues after DH6 recoding to account for possible compositional heterogeneity due to high GC% in cyclostome genomes. (**d**) Timetree inferred from the same dataset as (**b**) using a set of calibrations defined in Table S5, the CAT+GTR+G4 model and the CIR relaxed clock (with soft-bound) as implemented in Phylobayes.

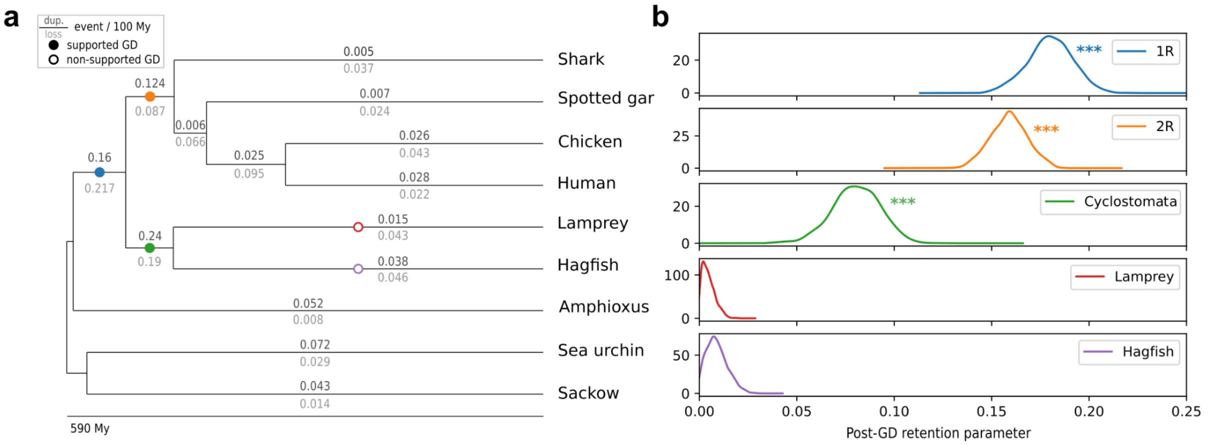


**Figure S4. Tests of genome duplication hypotheses on the vertebrate tree. (a)** Species phylogeny and Genome Duplication (GD) hypotheses. Genome duplication hypotheses are indicated by circles on the corresponding branches, with supported GD indicated as solid circles. Inferred background gene duplication and loss rates are presented on the branches. **(b)** Posterior distribution obtained for the WHALE post-GD retention parameters, for each hypothesis. Stars indicate distribution significantly different from 0, which corresponds to the supported GD events.

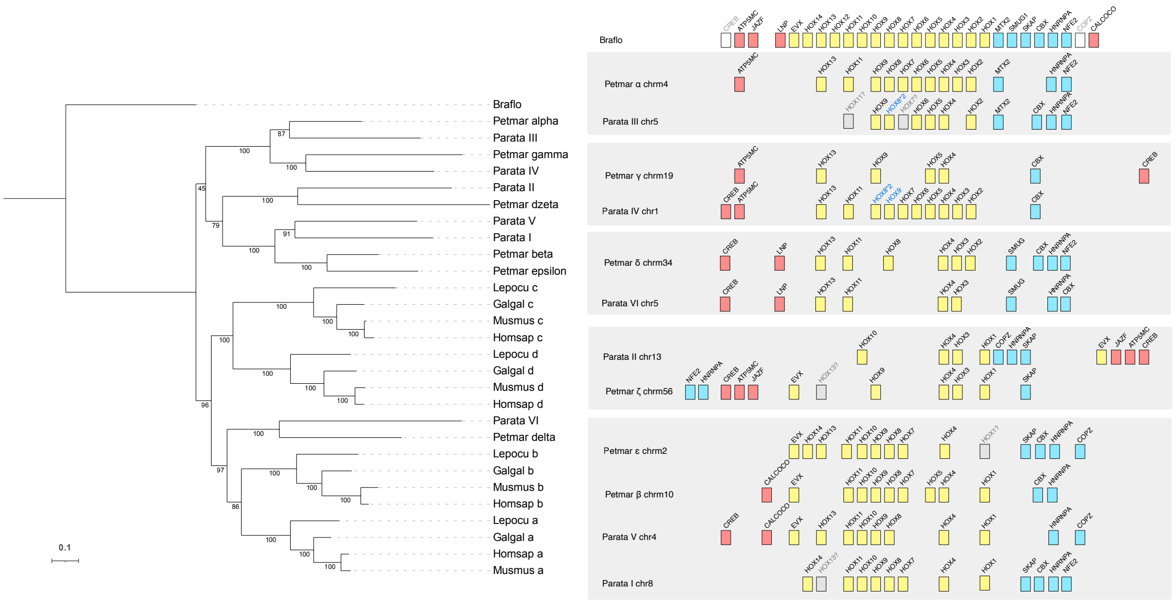


**Figure S5. Phylogeny of the Hox clusters based on a concatenation of Hox genes and bystanders. (a)** Phylogeny of the Hox clusters, with bootstrap support for nodes. One-to-one orthologies for gnathostomes clusters are well-supported, similarly for cyclostomes clusters with the exception of hagfish V/I and lamprey beta/epsilon. Dark gray boxes highlight cyclostomes clusters expected to be orthologs to gnathostomes clusters a/b based on chromosomal orthologies (**Figure 3b**), similarly light grey boxes are for expected orthologs to gnathostomes clusters c/d. **(b)** Schematic representation of cyclostomes and gnathostomes Hox clusters. Hox genes are shown as yellow boxes, 5’ bystanders as red boxes and 3’ bystanders as blue boxes. The order of genes reflects the actual arrangement of genes in each species.

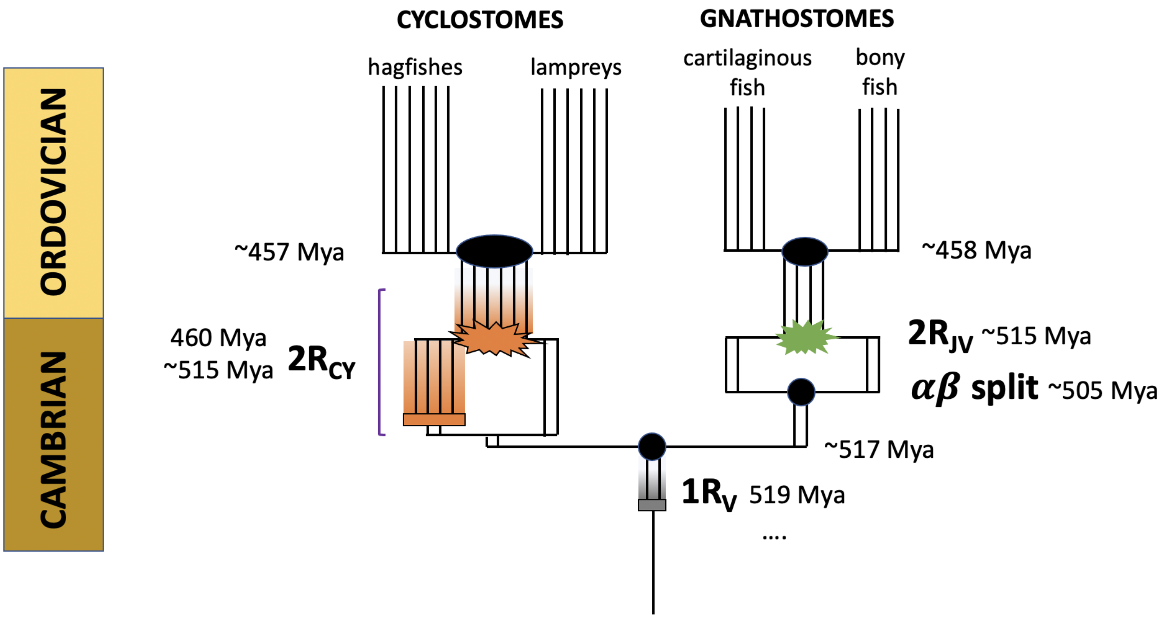


**Figure S6.** Scenario for genome duplication and speciation events during early vertebrate evolution. Filled black circles or ovals mark speciation events; coloured rectangles indicate presumptive autotetraploidies; starbursts indicate allopolyploidies arising from hybridization of distinct progenitors (e.g. alpha/beta in gnathostomes). Timings are based on **Figure 3c**. Note that while speciation times (e.g., the split between gnathostome progenitors alpha and beta) can be estimated from gene or paralogon trees, hybridization times (e.g., 2R$_{JV}$, shown as green starburst) cannot. The rough estimate of ~10 My interval between the alpha-beta split and 2R$_{JV}$ is based on analogy with recent vertebrate allotetraploidies in frogs and goldfish. Similarly, cyclostome hexaploidy (orange) is shown as arising from the hybridization of diploid and tetraploid stem cyclostomes, following the recent model of hexaploidy in sturgeon in which autotetraploids and diploid species coexist and hybridise. Although the cyclostome hexaploidy 2R$_{CY}$ is shown as two discrete events, the broad range of paralogon divergences in cyclostomes is consistent with an extended period of diploidization following autotetraploidy and/or hexaploidy arising from hybridization of closely related species that maintain homeologous recombination. Under these conditions, divergence times measure cessation of homeologous recombination, at which point loci can begin to evolve independently, a phenomenon noted by Furlong and Holland. This is shown as orange shading on the cyclostome stem. Similarly, the estimate of 1R$_V$ ~ 519 Mya represents the cessation of recombination after this presumptive autotetraploidy. (Autotetraploidy is suggested by the lack of differential gene loss between the two paralogous branches after 1R, as noted in Simakov et al.).

*Supplementary figures*

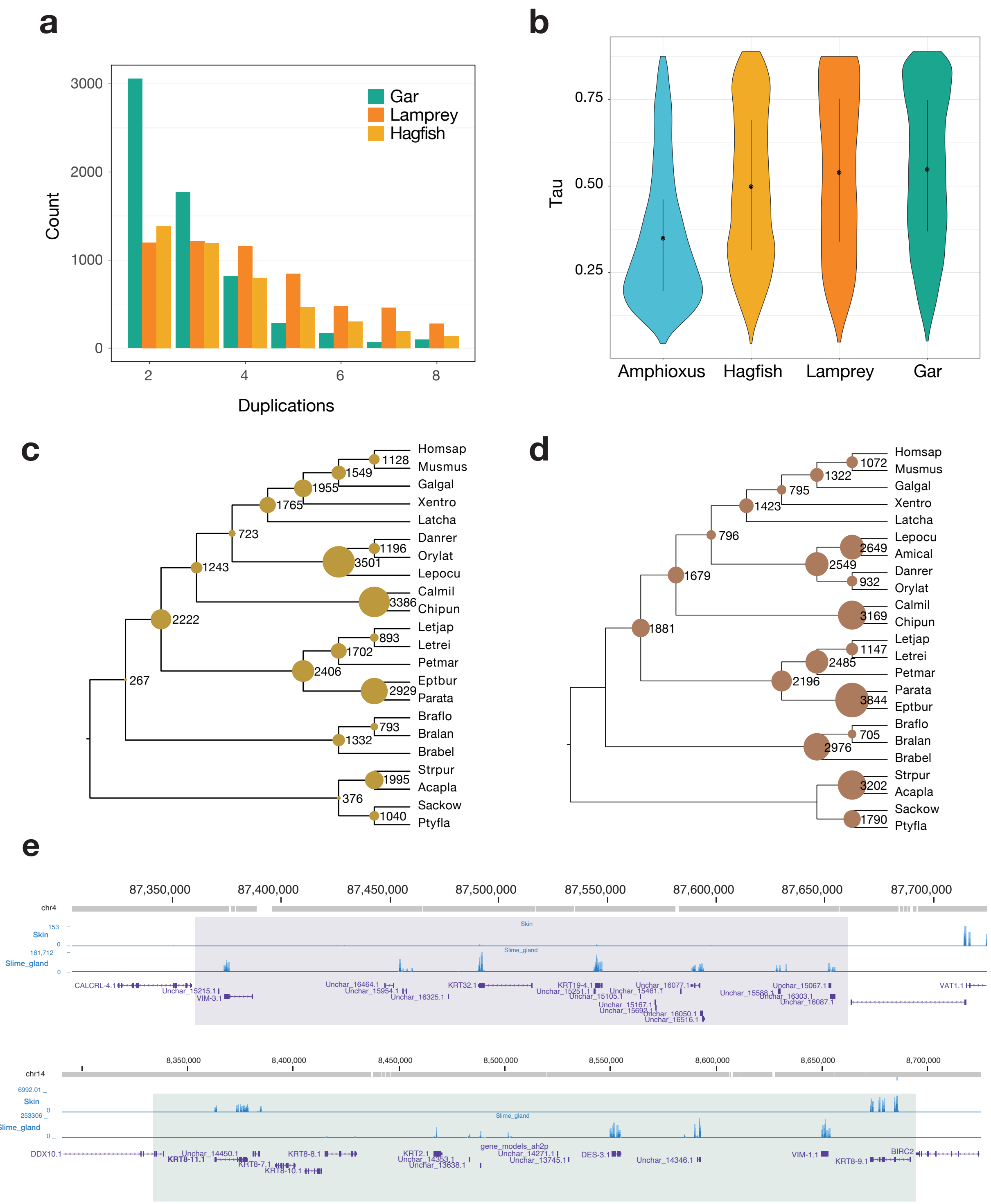


**Figure S7. Evolution of duplicated genes and gene families in hagfish** (**a**) Composition of paralogous gene families showing an ohnologue pattern in gar, lamprey (*P. marinus*) and hagfish (*E. atami*). (**b**) Comparison of the distribution of maximal tissue-specificity of gene expression (tau) in 6 tissues for members selected identical gene families in four chordates (**c**) Node-specific gene loss events as inferred by GeneRax in a species-gene tree reconciliation framework. (d) Loss of panther families across deuterostomes species, and (**d**) example of direct comparison of tau values between hagfish and amphioxus showing the maximal specificity for each paralogue in each species. (e) Genome structure of the two clusteds of expanded keratin genes.

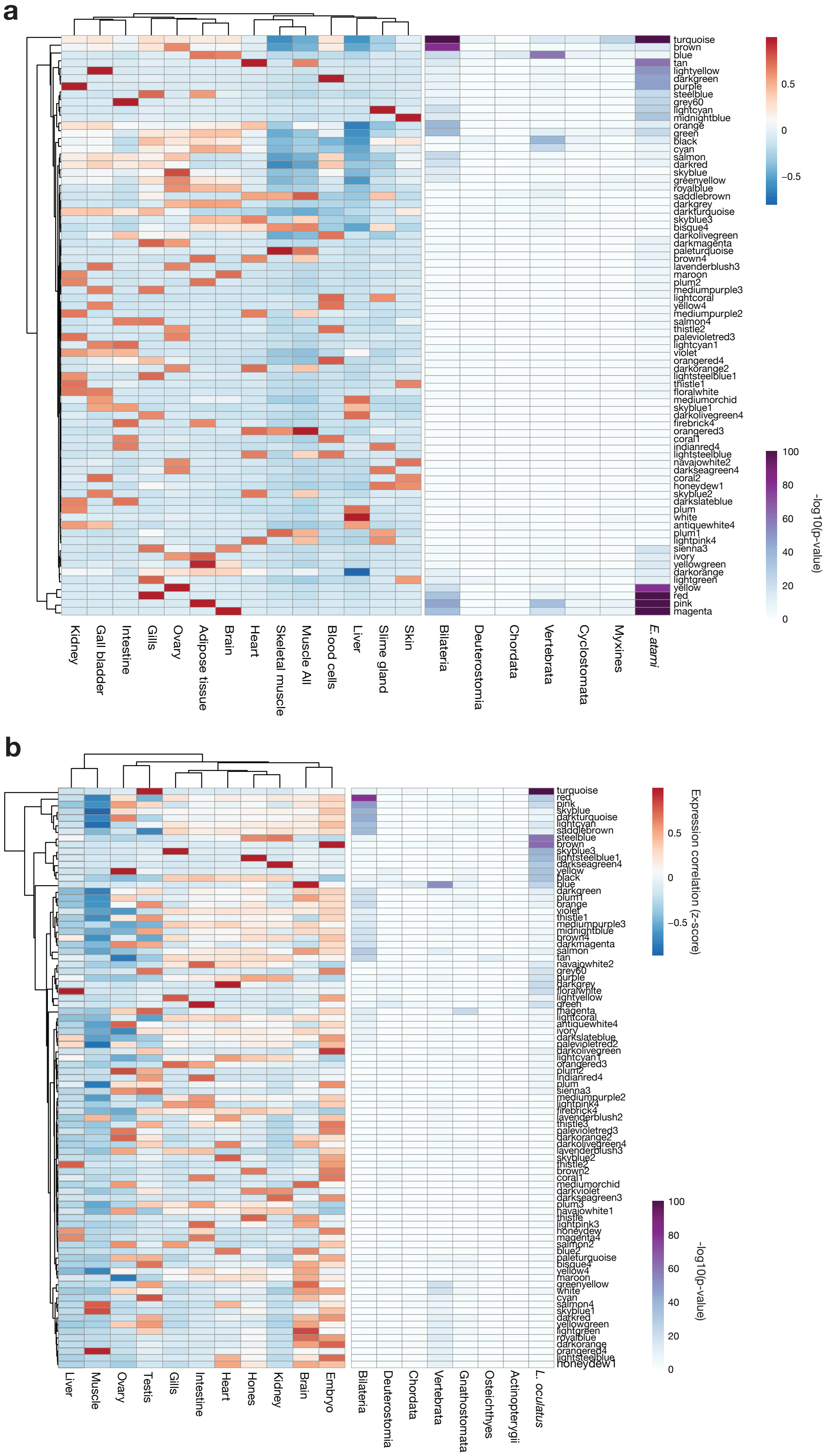


**Figure S8. Weighted gene co-expression network analysis (WGCNA) and gene duplications in vertebrates.** (**a,b**) On the left, WGCNA clusters obtained from gene expression in selected tissues of hagfish (a) and gar (**b**) and, on the right, hypergeometric enrichment of gene duplicated at successive phylogenetic nodes showing in both species.

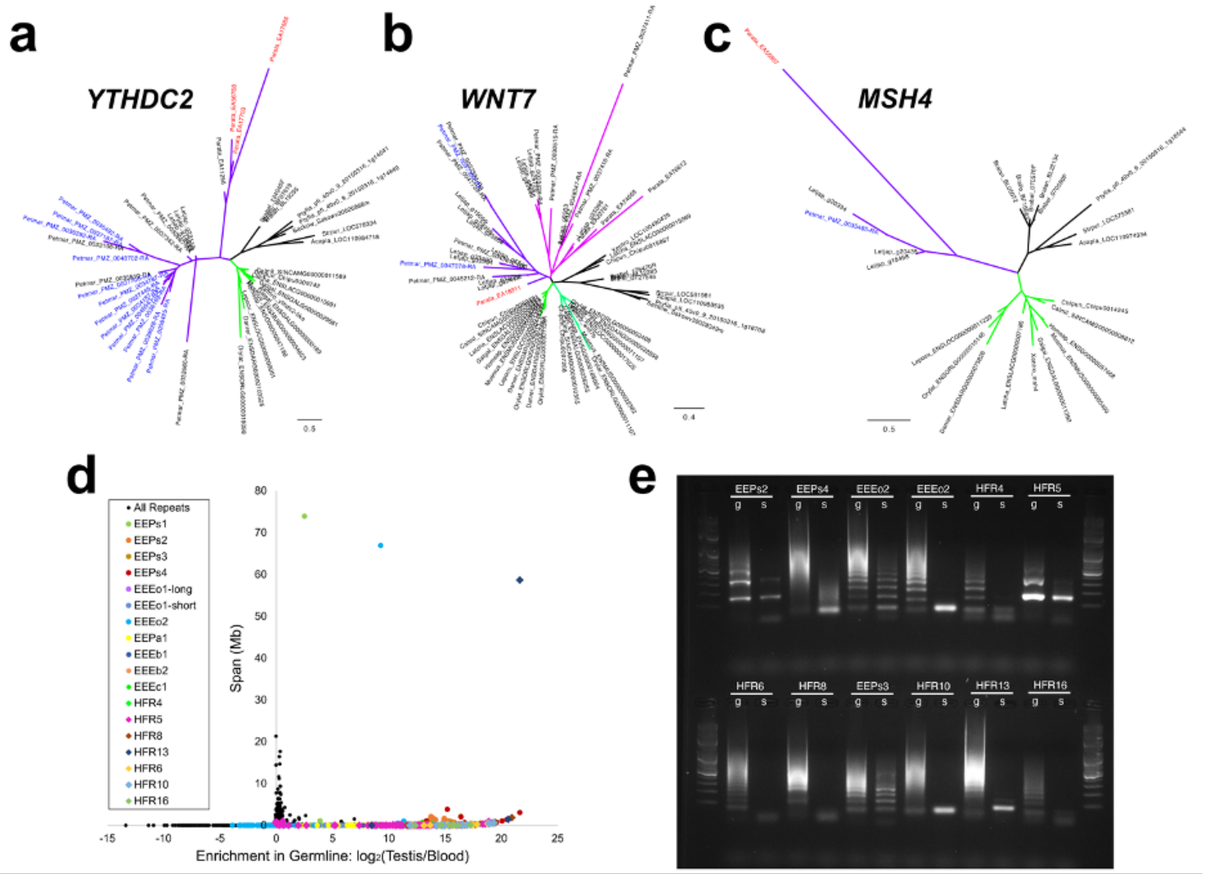


**Figure S9. Eliminated genes and repeats identified in the hagfish genome. (a-c)** Gene trees for homologs that are eliminated in both lamprey and hagfish. Gnathostome clades are highlighted in shades of green and cyclostome clades are highlighted in shades of purple. Individual germline-specific genes are highlighted in red (hagfish) or blue (lamprey). **(a)** Tree for YTHCD2 homologs. **(b)** Tree for WNT7 homologs. **(c)** Tree for MSH4 homologs. **(d)** Plot showing the degree of germline enrichment and estimated span of all predicted repetitive elements. Previously identified elements (Kojima et al., 2010; Kubota et al., 1993) are highlighted by coloured circles and new high copy elements are highlighted by coloured diamonds. **(e)** PCR validation illustrating germline enrichment and tandem repetition of predicted satellite elements. g: germline (testes) DNA used as template, s: somatic (blood) DNA used as template.

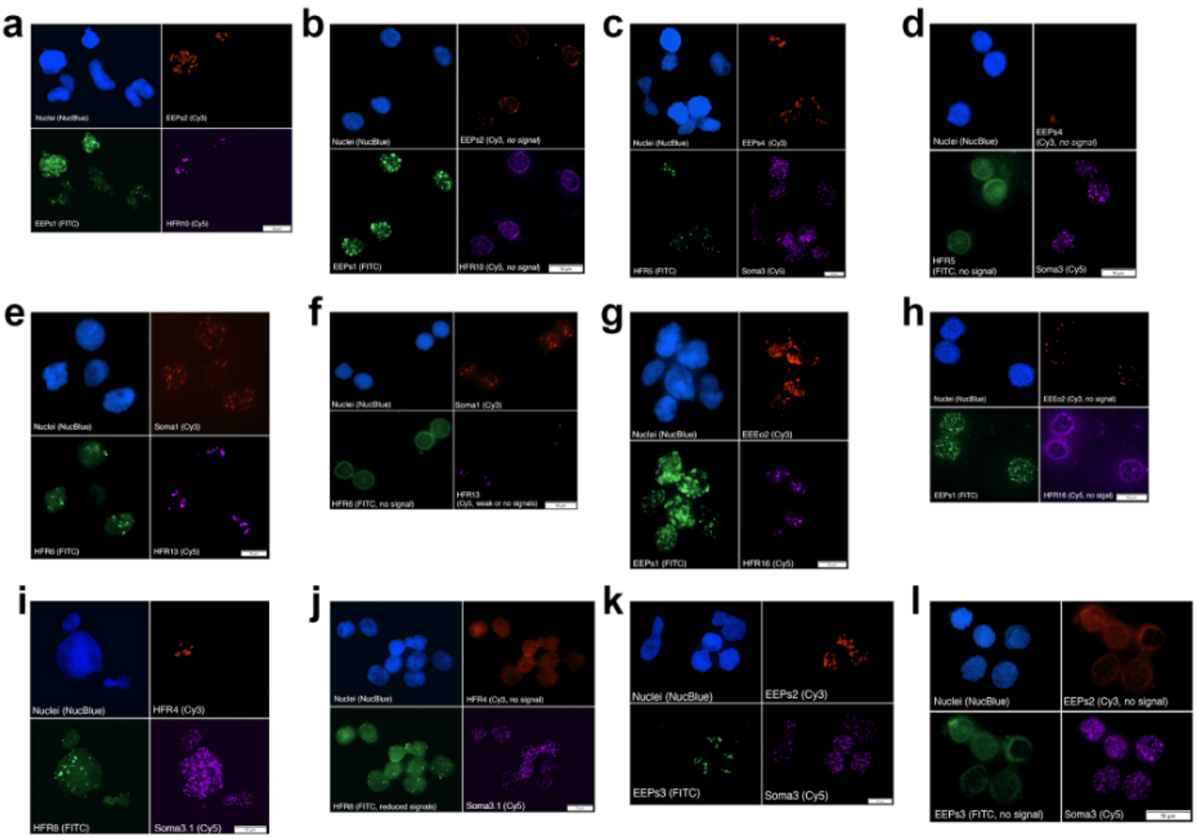


**Figure S10. Fluorescence in situ hybridization of repeats. (a&b)** Germline enriched EEPs2 (red) and HFR10 (magenta), and the somatic repeat EEPs1 (green) are hybridized to nuclei isolated from **(a)** germline: testes and **(b)** soma: blood. **(c&d)** Germline enriched repeats EEPs4 (red) and HFR5 (green), and the somatic repeat Soma3 (magenta) are hybridized to nuclei isolated from **(c)** germline: testes and **(d)** soma: blood. **(e&f)** Germline enriched repeats HFR13 (magenta) and HFR6 (green), and the somatic repeat Soma1 (red) are hybridized to nuclei isolated from **(e)** germline: testes and **(f)** soma: blood. **(g&h)** Germline enriched repeats EEEo2 (red) and HFR16 (magenta), and the somatic repeat EEPs1 (green) are hybridized to nuclei isolated from **(g)** germline: testes and **(h)** soma: blood. **(i&j)** Germline enriched repeats HFR4 (red) and HFR8 (green), and the somatic repeat Soma3.1 (magenta) are hybridized to nuclei isolated from **(i)** germline: testes and **(j)** soma: blood. **(k&l)** Germline enriched repeats EEPs2 (red) and EEPs3 (green), and the somatic repeat Soma3 (magenta) are hybridized to nuclei isolated from **(k)** germline: testes and **(l)** soma: blood.

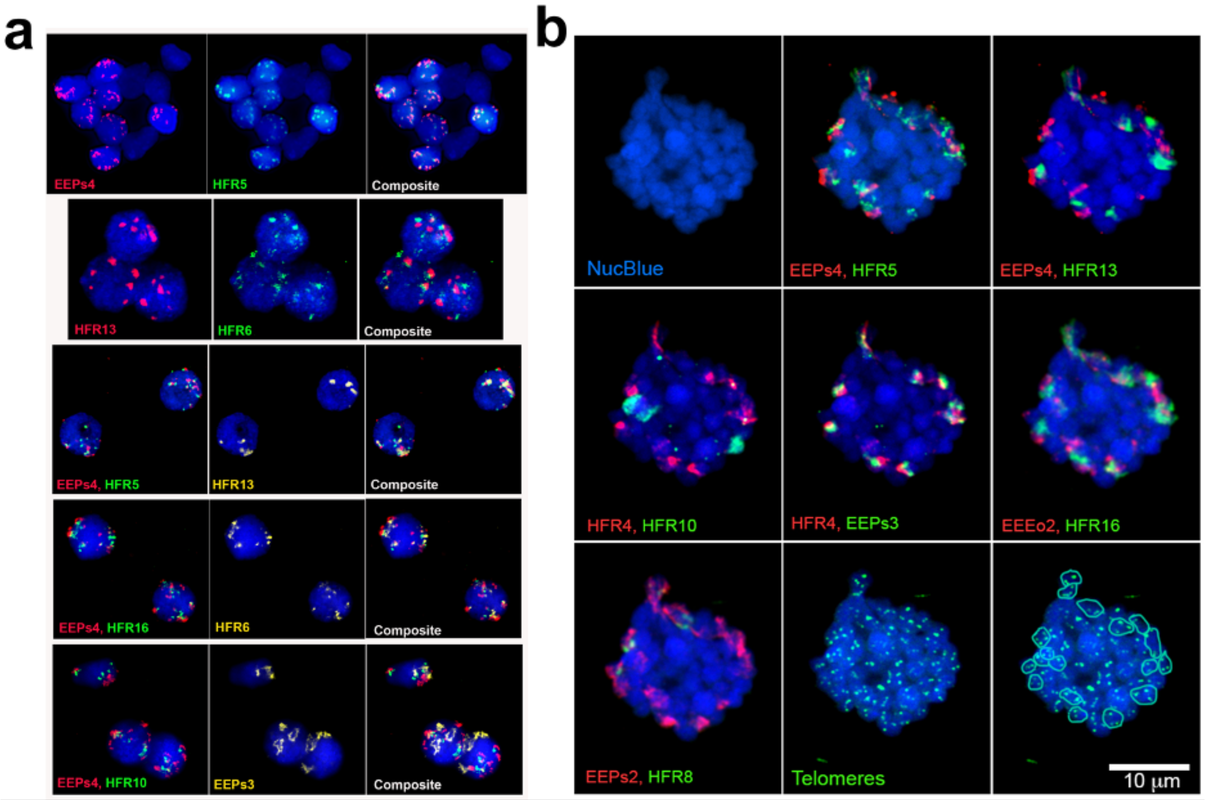


**Figure S11. In situ Hybridization of probes for ten germline-enriched satellite sequences. (a)** Probes are hybridized to germline interphase nuclei. **(b)** Probes are hybridized to germline interphase nuclei. The location of hybridization signals for telomere probes and approximate bounds of 18 germline-specific dyads, corresponding to nine distinct germline-specific chromosomes. For all images, pairs of repeats are shown to aid in visualizing the relative location of individual probes.
